# Genetic and preclinical evidence implicating chondroitin sulfate as a matritherapeutic target for the treatment of type 2 diabetes

**DOI:** 10.1101/2025.11.12.688143

**Authors:** Valdemar Brimnes Ingemann Johansen, Jens Lund, María José Romero-Lado, Alberte Wollesen Breum, Charlotte Svendsen, Kasper Suhr Jørgensen, Rebecca Louise Miller, Andreas Mæchel Fritzen, Tuomas O. Kilpeläinen, Katrine T. Schjoldager, Christoffer Clemmensen

**Affiliations:** Novo Nordisk Foundation Center for Basic Metabolic Research, Faculty of Health and Medical Sciences, University of Copenhagen, Copenhagen, Denmark; Copenhagen Center for Glycomics, Faculty of Health and Medical Sciences, University of Copenhagen, Copenhagen, Denmark; Department of Biomedical Sciences, Faculty of Health and Medical Sciences, University of Copenhagen, Copenhagen, Denmark

**Keywords:** Extracellular matrix, diabetes, type 2 diabetes, chondroitin sulfate

## Abstract

Peptide-based treatments for type 2 diabetes (T2D) are often limited by variable patient responses, frequent discontinuation, and substantial costs. Emerging lines of evidence link the extracellular matrix (ECM) to the pathophysiology of T2D, highlighting a largely unexplored modality for managing this heterogenous disease. Chondroitin sulfate (CS), a major glycosaminoglycan in the ECM, has been suggested to improve cardiometabolic health in preclinical research. However, the human genetic and pharmacological basis for CS as an anti-diabetic target is largely unexplored. Here, we uncover novel and robust links between 12 CS-related genes and both glycemic traits and the risk of T2D in humans. Complementing this, administration of CS leads to a profound lowering of blood glucose levels in severely diabetic mice and improves glucose tolerance and cardiac clearance of circulating glucose in diet-induced obese mice without causing hypoglycemia or other adverse effects. The improvement in glycemic control is accompanied by increased glucose-stimulated insulin secretion and enhanced insulin action, effects which seem to occur independent of the incretin system. The combination of human genetic evidence and appealing pharmacodynamic features highlights CS as a promising ECM-target for developing novel pharmacotherapies that complement current treatments for T2D.

## Introduction

The prevalence of diabetes has increased by more than 600 million people since the 1990s^1^ and is estimated to affect 1 in 8 adults worldwide by 2050^2^. The vast majority of individuals with diabetes suffer from type 2 diabetes (T2D)^3^, a polygenic and multifactorial disease characterized by hyperglycemia. When improperly controlled, elevated blood glucose leads to glucotoxic damages to blood vessels, eyes, kidneys, and nerves thereby driving microvascular complications and development of atherosclerotic cardiovascular disease^3^. The pathophysiology of T2D involves insulin resistance across multiple tissues, impaired pancreatic insulin secretion, and excessive hepatic glucose production^3–7^. However, other factors are emerging as additional drivers of the hyperglycemia, highlighting that the management of T2D, including the prevention of long-term cardiovascular complications^8,9^, might be improved by looking beyond the classical ‘ominous octet’.

While the ‘ominous octet’ has been instrumental in conceptualizing the pathophysiology of type 2 diabetes, it primarily emphasizes intracellular and organ-level defects that contribute to disease development or progression. Yet, this framework does not account for the extracellular matrix (ECM). The ECM is an essential and dynamic hydrogel meshwork of glycans and glycoconjugates in tissues. Here, it governs tissue architecture and signaling, as it constitutes a reservoir of growth factors and signaling molecules^10–12^. Interestingly, emerging evidence suggests that ECM remodeling may actively contribute to ý-cell failure and insulin resistance in skeletal muscle, the liver, and adipose tissue^13,14^. Elevated serum levels of ECM fragments have also been linked to increased risk of kidney complications in patients living with T2D^15^. Chondroitin sulfate (CS) is a major glycosaminoglycan found on proteoglycans in the ECM and is composed of negatively charged polysaccharide chains of repeating disaccharide units^16^. The biological functions of CS in health and disease, including cancer, inflammation, and fibrosis, are multifaceted and are largely determined by structural features, such as the CS chain length, its sulfation pattern, and the proteoglycan core^16^. Importantly, modulation of CS sulfation, epimerization, and structure has been implicated in the pathogenesis of diabetic nephropathy, retinopathy, and other aspects of diabetes complications^17,18^.

Over the last decade, ECM-based biologics and therapeutics have gained increasing attention as a promising new avenue to inspire novel pharmacotherapies, most notably in regenerative medicine^19,20^ and cancer^21,22^. These so-called matritherapies might also have potential for treatment of cardiometabolic diseases. In Europe and the United States, CS is approved and prescribed as a dietary supplement and a slow-acting drug for alleviating pain and other symptoms of osteoarthritis^23,24^. Interestingly, a few studies in streptozotocin-induced diabetes models have indicated that CS may improve blood glucose control in a type 1 diabetes-mimicked setting^25,26^, which raises the possibility that CS may also have therapeutic efficacy in preclinical models of T2D.

At present, it is uncertain whether the anti-diabetic effects of CS are applicable to humans, and whether CS may improve glucose tolerance in diet-induced obesity (DIO), a model with higher external and face validities in the context of T2D compared to streptozotocin-induced hyperglycemia^27^. Investigating if genetic variants affecting CS-related genes are associated with glycemic traits or T2D risk could uncover human evidence for CS as a matritherapeutic target in T2D and increase the likelihood of success in developing CS-based agents for managing hyperglycemia.

Here, we aimed to investigate the therapeutic potential of CS in T2D by screening CS-related human genes for associations with glycemic traits and T2D risk in the largest available genome-wide association studies (GWAS), and by investigating the effects of CS administration on glycemic control and energy balance in obese and diabetic mice.

## Results

### Human genetic evidence links chondroitin sulfate biology to glycemic traits and type 2 diabetes

Therapies targeting mechanisms that are supported by human genetics show improved rates of clinical success and are more likely to be approved^28,29^. To investigate whether genes encoding proteins involved in CS metabolism contribute to glycemic regulation in humans, we curated a panel of 28 genes implicated in CS metabolic pathways^30^: 13 genes involved in biosynthesis of CS, 9 in sulfation of CS, and 6 in degradation of CS (**Fig S1**, **Table S1**). We systematically screened these genes for associations with glycemic traits (fasting blood glucose, random blood glucose, and HbA_1c_ levels) and T2D risk using both single variant and gene-level approaches (**Fig 1A**). The screen revealed robust associations for 12 genes: seven involved in CS biosynthesis, two in CS sulfation, and three in CS degradation (**Fig 1A**).

**Figure 1.**
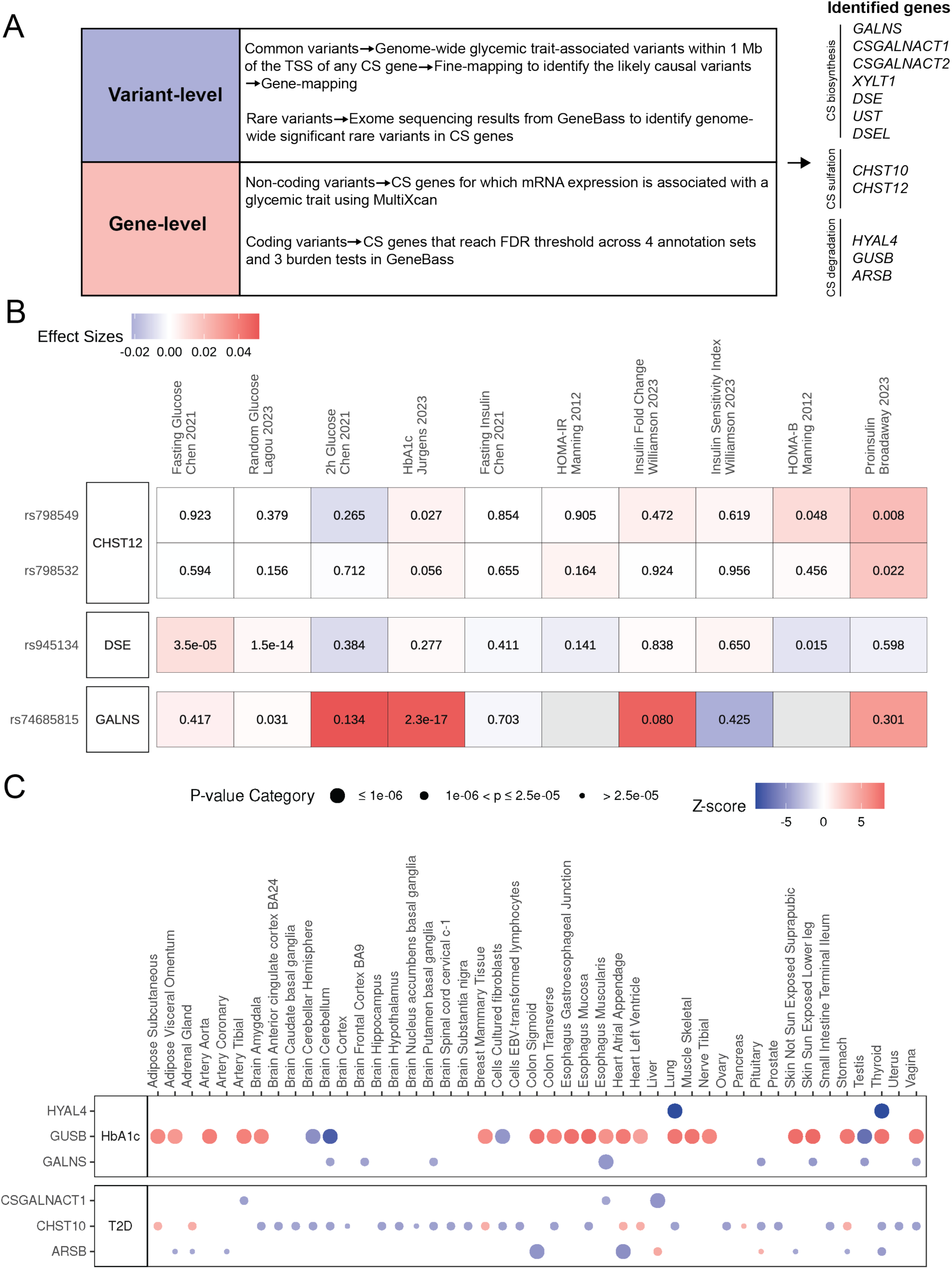
Human genetic screen for associations between CS genes and glycemic traits and T2D risk. **A)** Bioin-formatics workflow: Starting with 28 genes related to CS biosynthesis and degradation, we employed variant-level and gene-level approaches to identify genetic associations with T2D risk and glycemic traits (fasting glucose levels, random glucose levels, and HbA1c levels). In the variant-level analyses, we identified common variants through statistical fine-mapping and SNP-to-gene mapping approaches using the largest available GWAS for each trait. We also identified rare coding variants using the UK Biobank exome sequencing data. In the gene-level analysis, we investigated whether differential mRNA expression of CS genes across different tissues was associated with the glycemic traits of interest and used gene collapsing analysis to study gene-level associations for variants in the coding regions of the CS genes. **B)** Single variant associations for the identified common variants across several glycemic markers: The SNPs are on the y-axis and the traits are on the x-axis. The color indicates the effect sizes. Whenever possible the BMI-adjusted traits were used. **C)** Gene-level mRNA expression analysis for CS-related genes and glycemic traits: Six genes reached significance for one or more tissues within the GTEx Consortium data. The size of the dots indicates the level of statistical significance, and the color indicates the z-score. Only associations reaching the multiple testing threshold (P = 0.05/(49*28) = 3.64x10^-5^) are shown.

The seven genes found related to CS biosynthesis were: *GALNS*, *CSGALNACT1*, *CSGALNACT2, XYLT1*, *DSE*, *UST*, and *DSEL* (**Fig 1A**). The *GALNS* gene was identified through a common variant, rs74685815, associated with elevated HbA_1c_ levels (P=2x10^-17^), which was also nominally associated with random blood glucose levels (P < 0.05) (**Table 1**, **Fig 1B**). In addition, a rare synonymous variant Thr56Thr (Minor Allele Frequency, MAF_EUR_ = 0.01%) and gene-level evidence for both synonymous coding variants and mRNA expression further supported the role of *GALNS* in the regulation of HbA_1c_ levels (**Table 1**, **Fig 1C**). Pathogenic loss-of-function variants in *GALNS* are known to cause mucopolysaccharidosis type IVA, a rare congenital disorder characterized by e.g. growth retardation and skeletal abnormalities. Interestingly, a case report has documented co-occurrence of this condition with type 1 diabetes^31^. However, another and more severe case of mucopolysaccharidosis type IVA described normal glucose, insulin, and lipid profiles^32^.

**Table 1:**
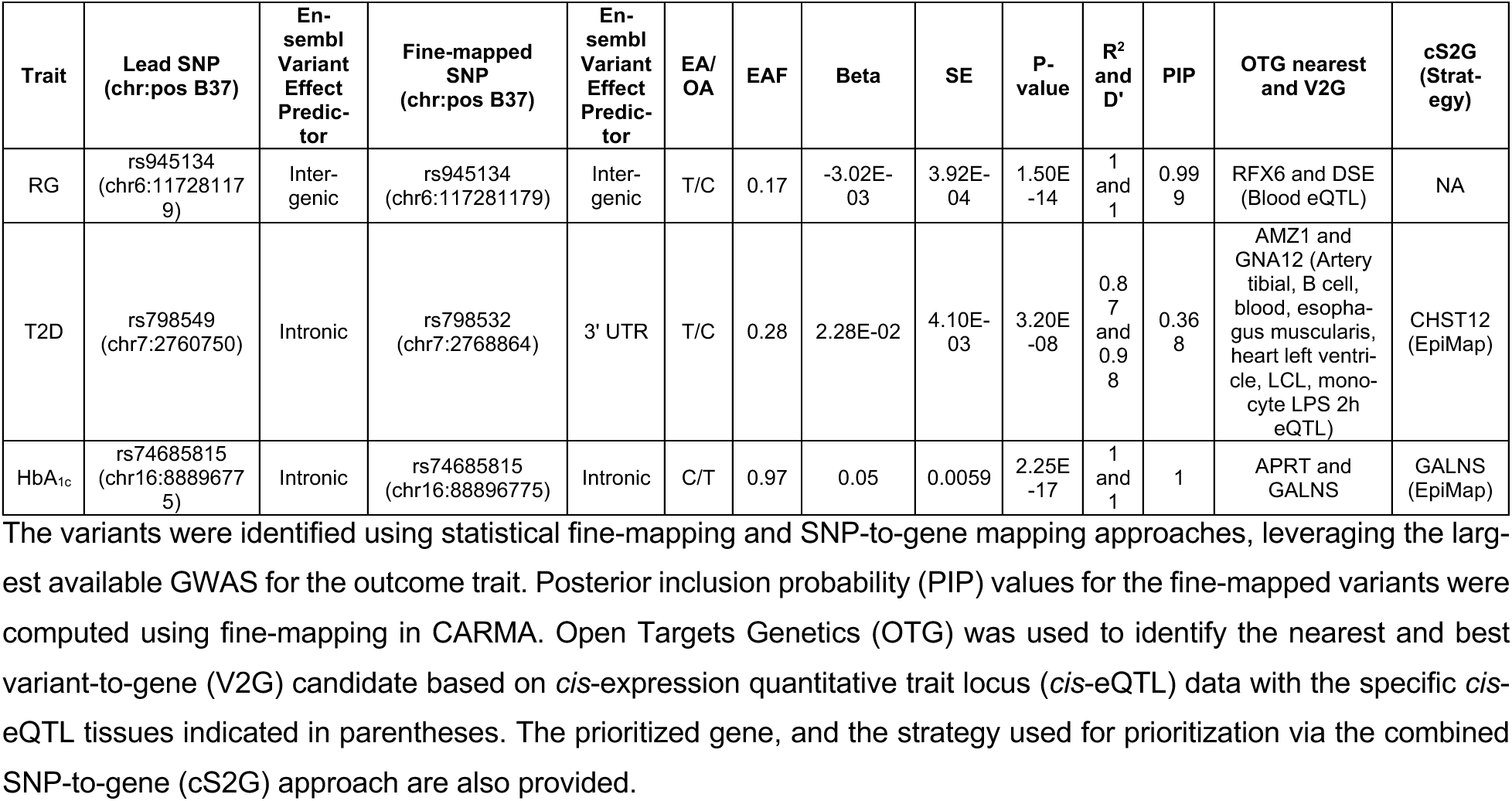
Common variants associated with glycemia or type 2 diabetes risk within regions defined by CS genes.

*CSGALNACT1* was associated with T2D risk through mRNA expression in the liver, esophagus, and tibial arteries (**Fig 1C**). The link between *CSGALNACT1* and T2D is further supported by the observation that *db^-^/db^-^* mice display lower endothelial expression of *Csgalnact1*^33^. *CSGALNACT2* was identified through predicted loss-of-function (pLoF) variants associated with insulin-dependent diabetes mellitus (ICD-10 code E10) (**Table 2**). Gene-level associations for *XYLT1* identified pLoF variants that are significantly associated with diabetes diagnosed by a physician, non-insulin-dependent diabetes mellitus (ICD-10 code E11), and unspecified diabetes mellitus (ICD-10 code E14) (**Table 2**). *DSE* was identified as the effector gene for the common variant rs945134 (MAF = 17%) associated with random blood glucose levels (P=1.5x10^-14^), and nominally with fasting blood glucose, random blood glucose, and HOMA-B (P < 0.05) (**Fig 1B**, **Table 1**). *UST* was associated through synonymous variants with other specified diabetes mellitus (ICD-10 code E13) (**Table 2**), while *DSEL* was associated with glucose through synonymous variants (**Table 2**).

**Table 2:** Gene-based association results for candidate glycemia-related CS genes.

| Gene symbol | Variant annotation | Genomic interval | Phenotype | Number of cases | Number of controls | Number of variants | Beta | Standard Error | Significant tests | Significant p-values |
| --- | --- | --- | --- | --- | --- | --- | --- | --- | --- | --- |
| CSGALNACT2 | pLoF | [chr10:43155108-43183596] | Date E10 first reported (insulin-dependent diabetes mellitus) | 3251 | 391590 | 38 | 0,09304 | 0,034507 | SKAT | 1.58e-03 |
| DSEL | synonymous | [chr18:67510906-67514695] | Glucose (NMR metabolite) | 94796 |  | 215 | -0,002643 | 0,00097288 | SKAT, SKAT-O | 1.36e-03, 1.74e-03 |
| GALNS | synonymous | [chr16:88809568-88861040] | Glycated haemoglobin (HbA<sub>1c</sub>) | 376931 |  | 168 | -0,0014994 | 0,00038101 | Burden, SKAT, SKAT-O | 8.31e-05, 1.28e-06, 1.70e-06 |
| UST | synonymous | [chr6:148747297-149074211] | Date E13 first reported (other specified diabetes mellitus) | 245 | 394596 | 45 | 0,15261 | 0,084792 | SKAT | 9.01e-04 |
| XYLT1 | pLoF | [chr16:17108602-17470893] | Diabetes diagnosed by doctor | 18980 | 374778 | 33 | 0,074867 | 0,023151 | Burden, SKAT-O | 1.22e-03, 9.70e-04 |
| XYLT1 | pLoF | [chr16:1710860<br>2-17470893] | Date E11 first reported (non-insulin-dependent diabetes mellitus) | 24153 | 370688 | 33 | 0,069<br>945 | 0,02170<br>5 | Burden,<br>SKAT-O | 1.27e-03,<br>7.01e-04 |
| XYLT1 | pLoF | [chr16:1710860<br>2-17470893] | Date E14 first reported (unspecified<br>diabetes mellitus) | 17397 | 377444 | 33 | 0,087<br>252 | 0,02269<br>6 | Burden,<br>SKAT, SKAT-O | 1.21e-04,<br>3.69e-04,<br>6.79e-05 |

This table summarizes significant gene–phenotype associations identified from GeneBass gene-level analyses after Bonferroni correction ( $\alpha = 0.05/28$ ). Variants are classified by functional annotation (predicted loss-of-function, pLoF or synonymous), and the associated glycemic phenotypes are provided using short clinical descriptions or biomarker traits. Effect size estimates (Beta and Standard Error) are shown. Columns “Significant tests” and “Significant p-values” indicate which statistical tests (Burden, SKAT, SKAT-O) passed the multiple testing threshold and the corresponding p-values.

The genes involved in CS sulfation and associated with glycemia and T2D risk include *CHST10* and *CHST12*. *CHST10* was identified through the association of mRNA expression with T2D risk across multiple tissues (**Fig 1C**). A common variant rs798532 (MAF = 28%) near *CHST12* was associated with T2D risk (**Table 1**, P = 3x10^-8^) and showed nominal association with proinsulin levels (P < 0.05) (**Fig 1B**). Furthermore, gene-level analysis of pLoF variants aligned with the association with diabetes mellitus in pregnancy (ICD-10 code O24) (SKAT-O P = 0.01184, SKAT P = 0.00794, burden P = 0.06178), although none of these P-values met the multiple-testing corrected threshold of 0.05/28 (0.00179).

The genes involved in the degradation of CS and showing robust associations with glycemic traits include *HYAL4*, *GUSB*, and *ARSB*. *HYAL4* and *GUSB* were both identified through associations between their mRNA expression and HbA_1c_ levels, with *HYAL4* showing tissue-specific associations in the lung and thyroid, and GUSB exhibiting associations across multiple tissues, including major blood vessels and the heart (**Fig 1C**). *ARSB* was identified as associated with T2D risk based on mRNA expression in multiple tissues (**Fig 1C**), and gene-level evidence for pLoF variants was consistent with the association with other specified diabetes mellitus (ICD-10 code E13) (SKAT-O P = 0.00503, SKAT P = 0.00335, burden P = 0.0457), although none of these P-values passed the multiple-testing corrected threshold of 0.05/28 (0.00179).

Taken together, our findings provide robust genetic evidence that distinct components of CS metabolism – biosynthesis, sulfation, and degradation – are linked to glycemic traits and T2D risk (**Fig 1**, **Table 1**, **Table 2**), highlighting the potential relevance of CS to gly-cemic regulation in humans.

### Chondroitin sulfate improves glycemic control in diet-induced obese mice

Pre-clinical studies have demonstrated that high-fat diet-feeding increases the expression of glycoproteins carrying CS within perineuronal nets of the hypothalamus^34^. Moreover, dysregulation of CS in perineuronal nets of the hypothalamus can impair hypothalamic insulin sensing and drive weight gain and development of diabetes in rodents^34^. Enzymatic degradation of the perineuronal nets using chondroitinase leads to weight loss, enhanced insulin signaling, and improved glucose homeostasis^34,35^. While these effects have been attributed to easier access to hypothalamic neurons for hormones like insulin, the beneficial effects on glycemic control could in principle also be partly driven by the chondroitinase-mediated liberation of CS moieties that subsequently might act on their own, potentially as signaling molecules similar to matrikines^36,37^. Given these observations and our own findings that several genes related to CS metabolism were linked to glycemic traits and risk for T2D in humans (**Fig 1**), we next administered pharmacological doses of exogenous CS to mice to further explore the potential role of CS in glycemic control. Because of its low cost and use in human dietary supplements^32-35^, we used CS isolated from shark. This decision was further supported by the observation that orally administered CS from shark, but not from bovine cartilage, has been shown to lower the postprandial surge in blood glucose in streptozotocin-induced diabetic mice^38^.

Initially, we sought to determine if the glucose tolerance of DIO mice was affected by CS treatment. We examined the blood glucose excursions following intraperitoneal administration of glucose in combination with CS or vehicle administered subcutaneously 2, 3, 4, 5, and 6 h before the glucose dosing. As with CS treatment 30 min before intraperitoneal glucose (**Fig S2A-B**), CS dosed 2 h before did not affect glucose tolerance (**Fig 2A-B)** However, CS treatment 3, 4, 5, and 6 h before intraperitoneal glucose significantly improved glucose tolerance in DIO mice, where 6 h pre-treatment with CS numerically improved glucose tolerance the most (**Fig 2A-B**). An improvement in glucose tolerance was not observed in animals dosed with CS 21 h before the intraperitoneal glucose injection (**Fig S2C-D**).

**Figure 2.**
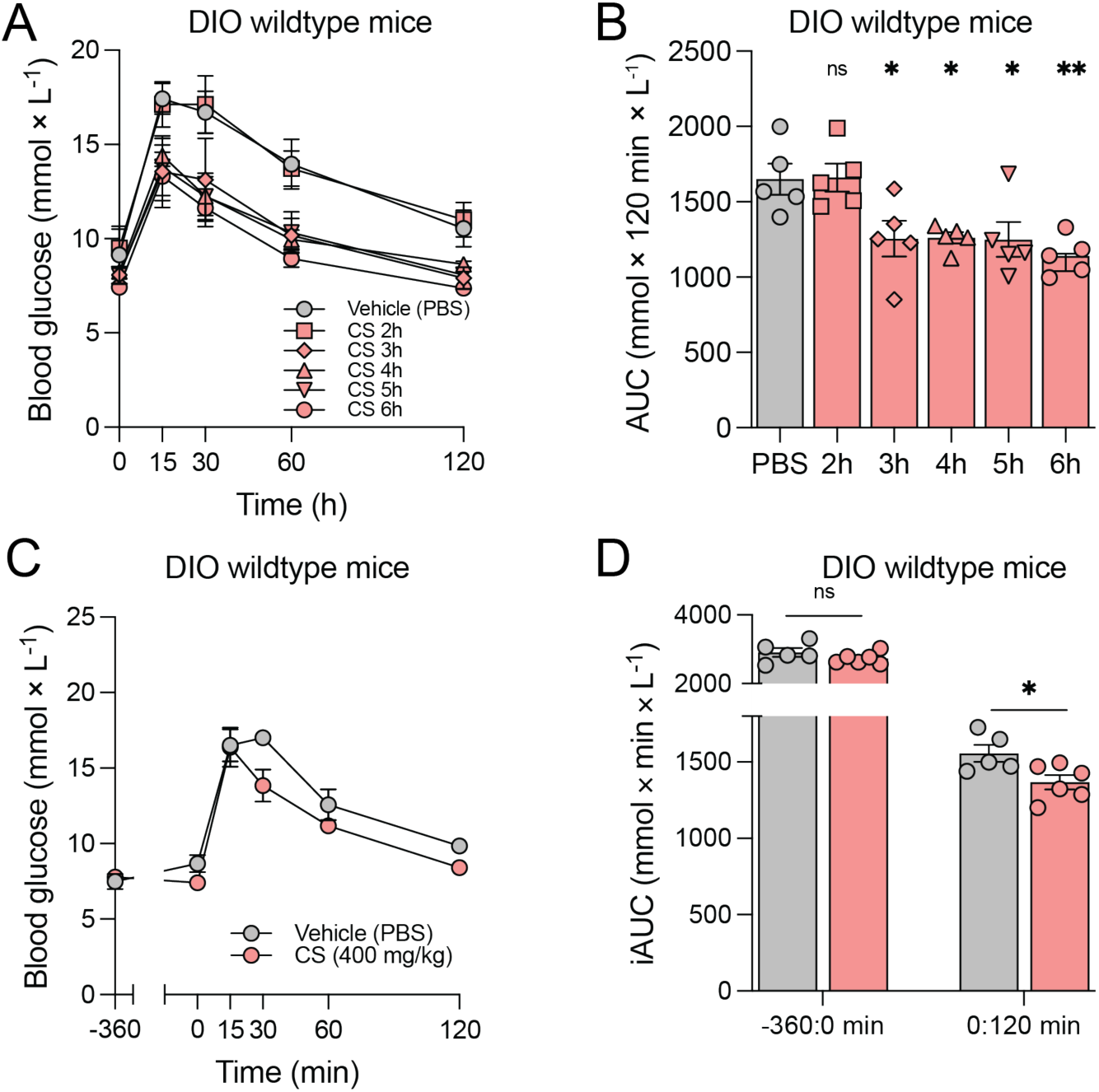
CS improves glycemic control in obese mice. **A-B)** Glucose tolerance tests in diet-induced obese (DIO) mice with time dependency of CS dosing. DIO mice were subcutaneously injected with vehicle 6 h (grey) or CS either 2, 3, 4, 5, or 6 h (red squares, rhombi, triangles pointing up, triangles pointing down, and circles, respectively, and as indicated) before an intraperitoneal injection of glucose, and their blood glucose levels were measured (panel C) and the AUCs calculated (panel D) (*n* = 5 per group) (mouse study 1 in Materials and Methods). **C-D)** DIO mice were subcutaneously injected with vehicle (grey) or CS (red) 6 h before an intraperitoneal injection of glucose, and their blood glucose levels were measured (panel E) and the iAUCs calculated, as indicated (panel F) (*n* = 5-6 per group) (mouse study 2 in Materials and Methods). Statistics in panels B and F by unpaired t-tests with Welch’s correction. Statistics in panel D by multiple unpaired t-tests with Welch’s correction against vehicle and correction for multiple testing. *, *p* < 0.05; **, *p* < 0.01; ns, not significant.

Given that we observed a modest, yet insignificant, decrease in blood glucose levels in DIO mice treated with CS compared to vehicle already before intraperitoneal glucose (**Fig 2A**, timepoint 0), we controlled for differences in basal blood glucose levels at the time of CS and vehicle treatment. This experiment replicated the finding of significantly improved glucose tolerance in DIO mice administered CS 6 h before intraperitoneal glucose (**Fig 2C-D**, 0:120 min), and we found that basal blood glucose levels were insignificantly decreased by CS compared to vehicle treatment (**Fig 2D**, -360:0 min). Consistent with these observations, 2, 4, and 6 h after acute CS treatment, basal blood glucose levels in DIO mice were unaffected by CS treatment compared to a vehicle treatment (**Fig S2E-F**). To control for a potential confounding effect arising from the viscosity of the CS solution, we administered viscous dextran and mannitol and found that this did not affect glucose tolerance in DIO mice (**Fig S3A-B**). Furthermore, neither oral (**Fig S3C-D**) nor intracerebroventricular (**Fig S3E-F**) CS administration improved glucose tolerance, suggesting that CS might exert its anti-diabetic effects through a peripheral rather than a central mechanism.

To understand if the observed CS-induced improved glucose tolerance in DIO mice was replicated in lean mice, we performed a glucose tolerance test following subcutaneous injection of CS or vehicle in lean mice fed a chow diet. We observed that treatment did not affect glucose tolerance in this model (**Fig S3G-H**) and that basal blood glucose levels were not affected by acute CS vs. vehicle dosing (**Fig S3I-J**). In sum, CS improves glucose tolerance in DIO mice, but not in lean mice. This indicates not only that CS might work specifically in individuals with impaired glycemic control, but also that the efficacy of CS somehow increases as the hyperglycemia worsens.

### Chondroitin sulfate increases glucose-stimulated, but not basal, insulin secretion and enhances insulin-stimulated glucose clearance

Our findings that CS treatment led to improved glucose tolerance in DIO mice (**Fig 2A-D**) and that CS had no acute effects on blood glucose levels in DIO mice (**Fig 2C-D**, **Fig S2E-F**) suggested that an increase in glucose-stimulated insulin secretion could drive the CS-dependent improvement in glycemic control in DIO mice. To evaluate this hypothesis, we exposed CS and vehicle-dosed DIO mice to an intraperitoneal injection of glucose, replicating the CS-dependent improved glucose tolerance (**Fig 3A-B**). We observed that CS-dosed animals responded to the glucose injection by a ∼10 % larger increase in plasma insulin levels compared to vehicle-treated mice (**Fig 3C-D**, 0:30 min). Plasma insulin fold changes from the treatment timepoint (-360 min) to the glucose injection timepoint (0 min), however, were not affected by the treatment (**Fig 3C-D**, -360:0 min). Also at time points 2, 4, and 6 h after acute CS treatment, plasma insulin levels were unaffected by CS compared to a vehicle treatment in DIO mice (**Fig 3E-F**) and mice fed chow (**Fig S4A-B**).

**Figure 3.**
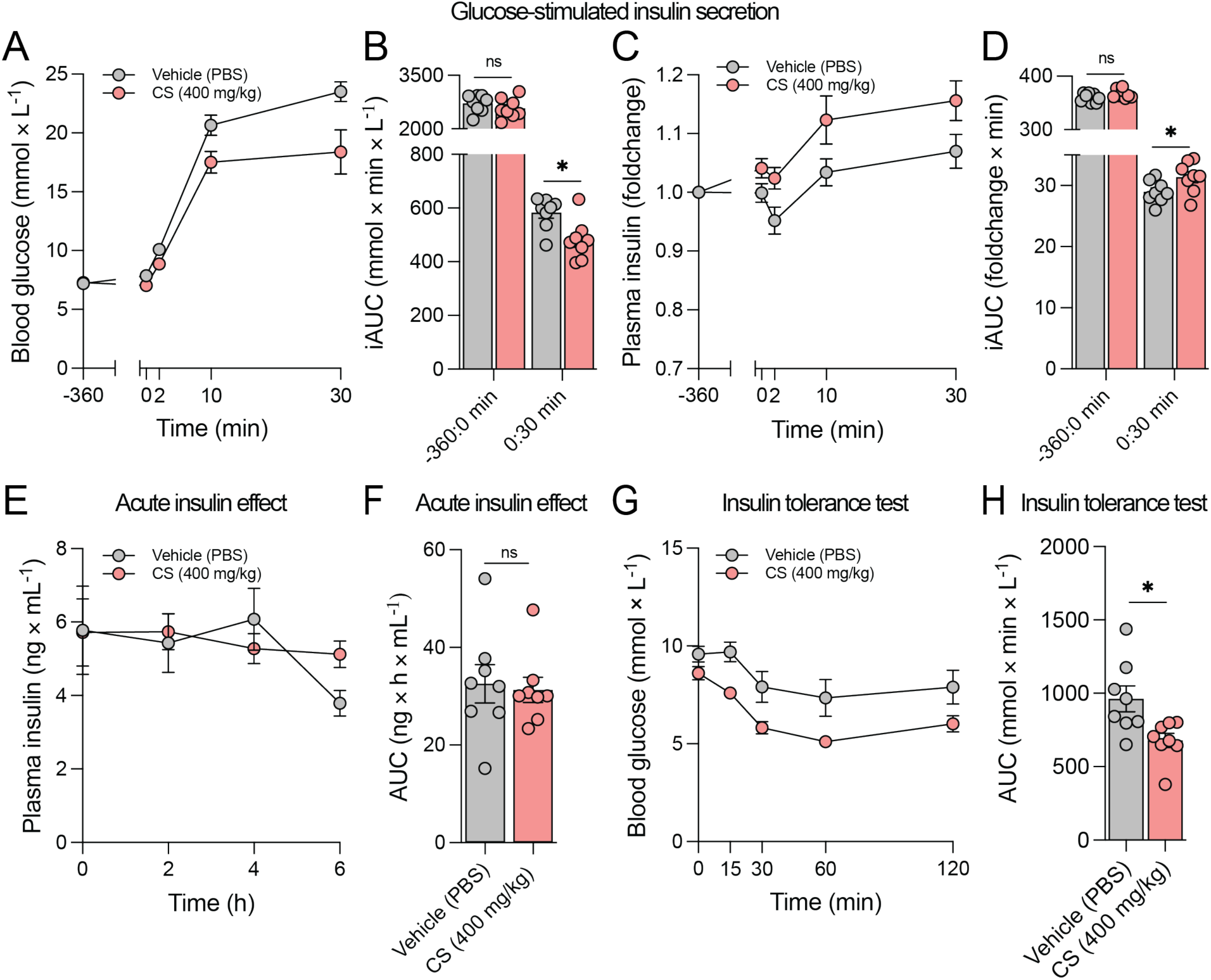
CS increases glucose-stimulated insulin secretion in diet-induced obese mice. **A-D)** Glucose stimulated insulin secretion in DIO mice dependent on vehicle (grey) or CS (red) treatment 6 h before an intraperitoneal glucose injection (*n* = 8 per group) (mouse study 3 in Materials and Methods). Blood glucose levels (panel A), blood glucose iAUCs (panel B), plasma insulin foldchange of baseline (panel C), and its iAUCs (panel D) from the experiment. **E-F)** Acute compound tolerance test of CS (red) and vehicle (grey) on plasma insulin levels in DIO mice (*n* = 8 per group) (mouse study 4 in Materials and Methods). **G-H)** Insulin tolerance test in DIO mice dependent on vehicle (grey) or CS (red) treatment 4 h before an intraperitoneal injection of insulin (*n* = 8 per group) with blood glucose levels measured (panel G) and area under the curve (panel H) (mouse study 5 in Materials and Methods). Statistics in panels B, D, F, and H by multiple unpaired t-tests with Welch’s correction. *, *p* < 0.05.

To investigate if insulin-stimulated glucose clearance was affected by CS treatment, we injected DIO mice subcutaneously with CS or vehicle 4 h before an intraperitoneal insulin injection. CS compared to vehicle treatment led to a significantly increased insulin-stimulated glucose disposal (**Fig 3G-H**).

In sum, these data indicate that CS treatment increases glucose-stimulated insulin secretion and potentiates insulin action in DIO mice. The increased glucose-stimulated insulin secretion might be a major explanation for the improved glucose tolerance in DIO mice.

### Chondroitin sulfate increases glucose-stimulated glucose clearance in the heart

To investigate if the improved glucose tolerance in DIO mice treated with CS could be explained by an enhanced glucose clearance from the blood into tissues, we conducted a 2-deoxyglucose tracing study under glucose-stimulating conditions in DIO mice (**Fig 4**). This experiment replicated findings suggesting an improved glucose tolerance upon CS treatment in DIO mice with no effects on basal blood glucose levels (**Fig 4A-B**), and 2-deoxyglucose tracer levels in the blood were lower in CS compared to vehicle-treated mice (**Fig S5A-B**). This suggests, as expected, that clearance of the tracer was affected by CS treatment. Surprisingly, the heart displayed significantly increased glucose-induced glucose clearance upon CS compared to vehicle treatment, while glucose clearance by other tissues was unaffected by treatment (**Fig 4C**). The enhanced glucose-induced glucose clearance into cardiac tissue in CS-treated mice prompted us to explore if this was associated with induction of insulin signaling in the heart. By western blotting of cardiac tissues from vehicle and CS-treated mice, we found no detectable differences in phosphorylated protein levels of pTBCD4^642^, pAKT308, and pAKT^473^, three key phospho-proteins that mediate intracellular signaling cascades downstream of insulin receptor activation (**Fig 4D-E**). Taken together, these data suggest that the improved glucose tolerance by CS compared to vehicle treatment in DIO mice is accompanied by increased glucose-induced glucose clearance into the heart, while these differences are not reflective of changes in the evaluated insulin signaling components dependent on treatment. It remains possible that the increased glucose clearance by the heart is mediated by other factors such as increased cardiac perfusion, insulin-induced signaling molecules not assessed here, or via insulin-independent mechanisms.

**Figure 4.**
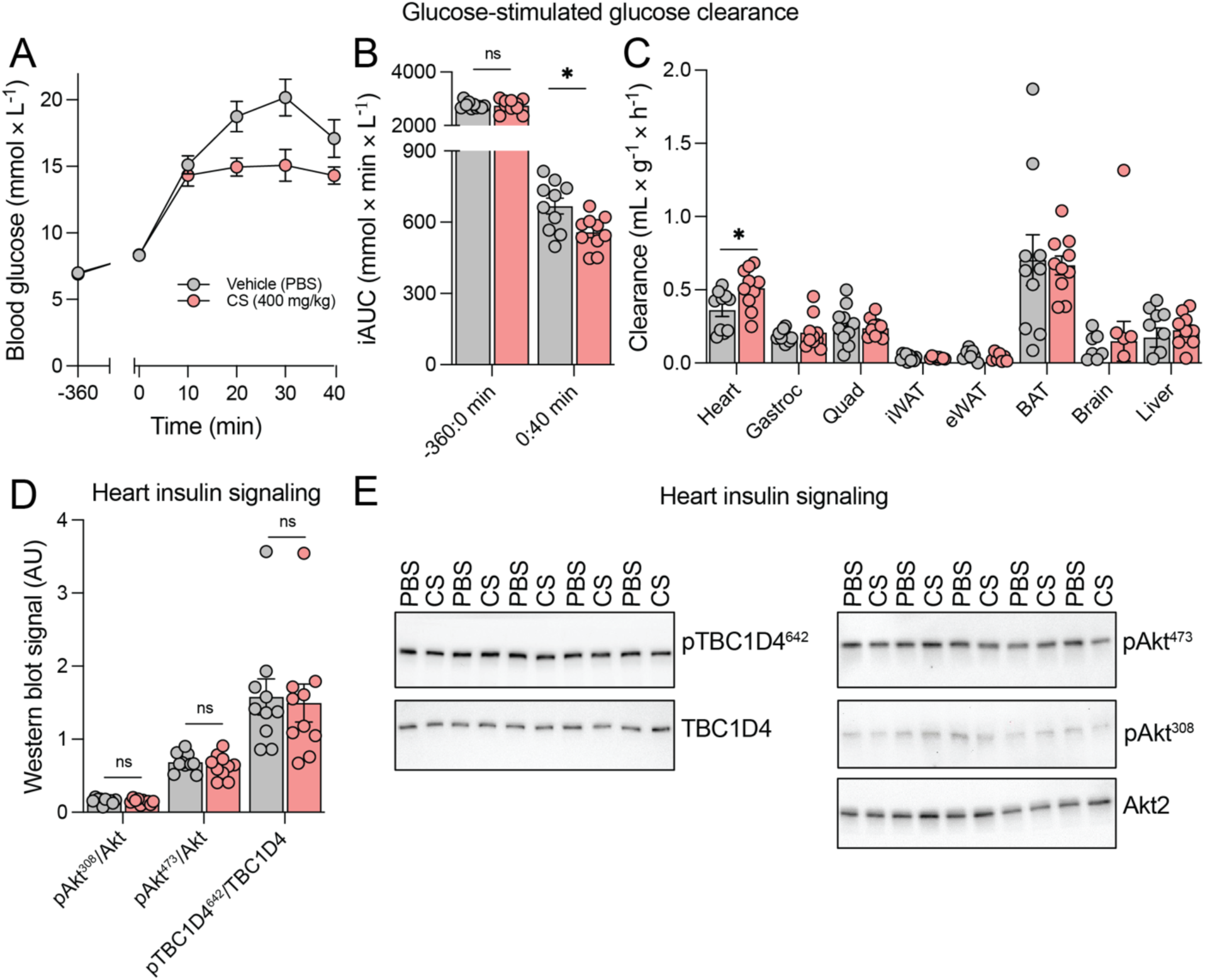
CS treatment increases glucose clearance from the blood into the heart under glucose-stimulating conditions. **A-C)** Glucose-stimulated glucose clearance dependent on vehicle (grey) or CS (red) treatment (*n* = 10 per group) 6 h before an intraperitoneal injection of glucose with trace amounts of ^3^H-2-deoxyglucose (mouse study 6 in Materials and Methods). Blood glucose levels (panel A), its incremental areas under the curve (iAUC) (panel B) and glucose clearance of indicated tissues (panel C). **D-E)** Quantified western blot signals (panel D) and representative western blots (panel E) of indicated insulin signaling components of hearts from DIO mice dosed with vehicle or CS. Statistics in panels B, C, and D by multiple unpaired t-tests with Welch’s correction. *, *p* < 0.05.

### Pharmacological chondroitin sulfate does not trigger aversion or sickness behaviors

Intrigued by the improved control of blood glucose levels by CS treatment in DIO mice (**Fig 2-4**), we investigated whether these beneficial effects of CS were associated with any adverse effects. It has previously been reported that physiological responses to various biologically active substances that take several hours to develop can be driven by cytokines that are released in response to biocontaminants like endotoxins^39,40^. This is also known as an ‘endotoxic reaction’ and can lead to e.g. fever and decreased blood glucose^41^. Moreover, the commercially available CS powder used here is a salt that contains sodium, a counterion that significantly increases the tonicity of the experimental solutions and therefore might induce malaise when injected subcutaneously to mice at high doses^42^. Physiological confounding from such biocontaminants is rarely considered in pre-clinical research^39,42^ and previous CS studies have not investigated whether the reported beneficial cardiometabolic effects of CS might be driven by adverse confounding from biocontaminants. We conducted a series of behavioral experiments in mice, including open field tests, voluntary running, and conditioned taste aversion (**Fig 5**). Compared to a vehicle injection, CS did not affect the behavior of mice in an open field (**Fig 5A-D, Fig S6A-E**).

**Figure 5.**
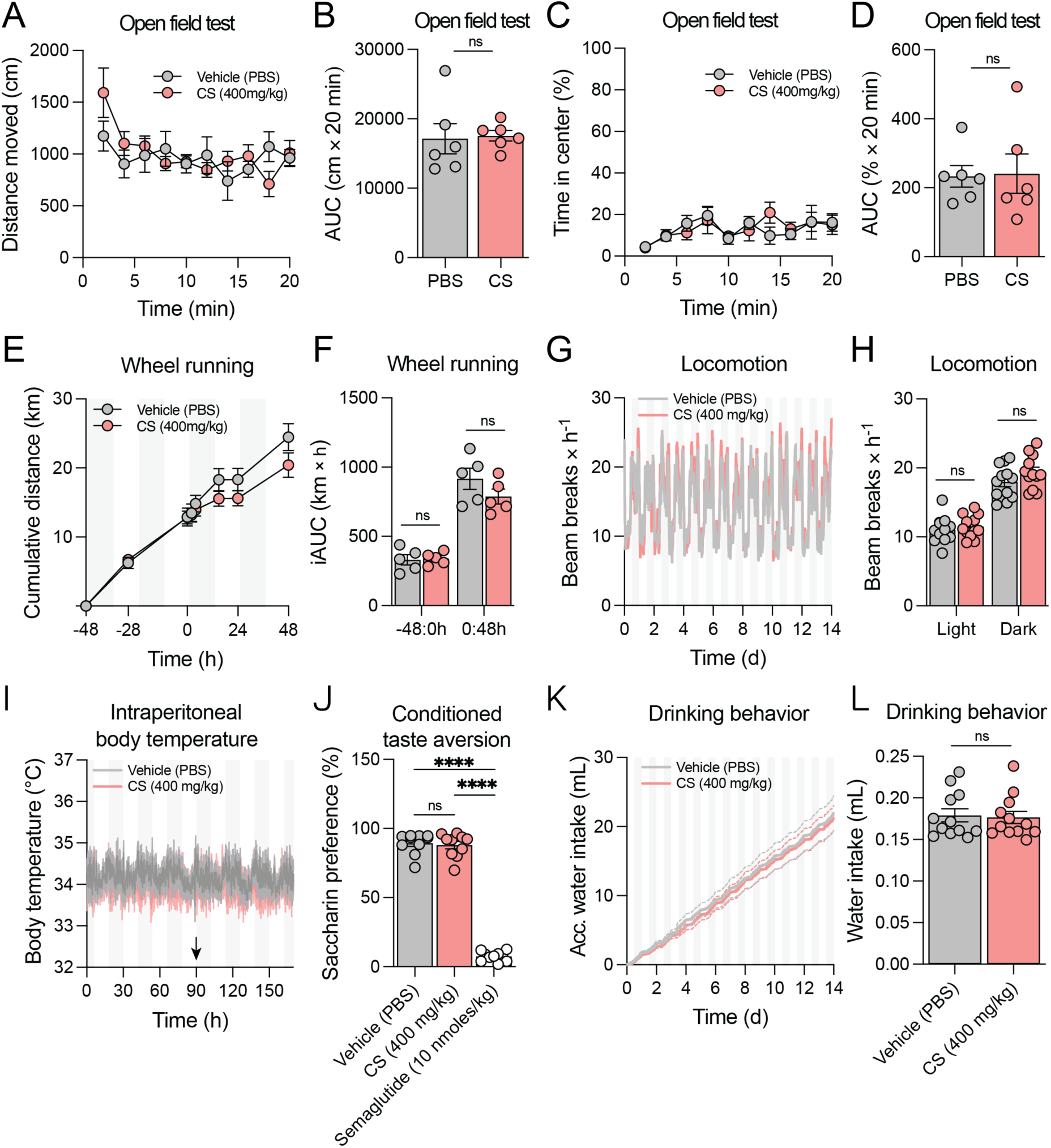
Behavioral assessments of acute CS administration *in vivo*. **A-D)** Open field test. Chow mice were subcutaneously injected with either CS (red) or vehicle (grey) (PBS) (*n* = 6 per group) and their behavior was followed acutely in an open field for 20 min. Distance moved (panel B) and its AUC (panel B), time spent in center (panel C) and its AUC were recorded or measured in intervals as indicated (mouse study 7 in Materials and Methods). **E-F)** Voluntary wheel running test. Chow mice were subcutaneously injected with either CS or vehicle (*n* = 5 per group) and their cumulative running distance (panel E) and its iAUCs (panel F) were measured 48 h before and after injection (mouse study 8 in Materials and Methods). **G-H)** Locomotor activity. DIO mice were subcutaneously injected daily with either CS or vehicle (*n* = 12 per group) in metabolic cages and their locomotion recorded (panel G) and average locomotor activity were calculated based on light or dark phases, as indicated (panel H) (mouse study 9 in Materials and Methods). **I)** Assessment of intraperitoneal body temperature. Chow mice were subcutaneously injected with either CS or vehicle (*n* = 5 per group) and their intraperitoneal body temperatures were measured before and after injection (mouse study 10 in Materials and Methods). Arrow indicates the time of injection. **J)** Conditioned taste aversion. Chow mice were subcutaneously injected with either CS, vehicle, or semaglutide (*n* = 10 per group) (mouse study 11 in Materials and Methods) with only access to saccharin (conditioning day). Twenty-four hours post-conditioning day, mice had free access to water and saccharin, and their saccharin preference was measured throughout 24 h. **K-L)** Drinking behavior. DIO mice were subcutaneously injected daily with either CS or vehicle (*n* = 12 per group) in metabolic cages and their drinking behavior recorded (panel K) and average water intake were calculated, as indicated (panel L) (mouse study 9 in Materials and Methods). Statistics in panels B, D, F, H, J, and L by multiple unpaired t-tests with Welch’s correction. Data presented as means +/-SEM and individual data points represent data from individual animals. ****, *p* < 0.0001; ns, not significant.

Further, we found no significant differences in the cumulative running distance and time from a voluntary running experiment, although a tendency of decreased running distance and time was observed in CS compared to vehicle-treated mice (**Fig 5E-F**, **Fig S6F-G**). In contrast, other drugs with aversive effects, like semaglutide, show prominent reductions in voluntary wheel running^43^. Consistent with the open field recordings, CS did not induce changes in locomotor activity in DIO mice treated daily for 14 days compared to vehicle-treated mice (**Fig 5G-H**, **Fig S6H-I**). This lack of locomotor effects from CS treatment was supported by results from another cohort of DIO mice (**Fig S7C**).

In line with the lack of any abnormal behavior, a single treatment with CS did not change the body temperature of mice (**Fig 5I**) and neither did CS acutely change the energy expenditure or respiratory exchange ratio of mice (**Fig S7A-B**). No evidence of conditioned taste aversion was observed upon injection of CS compared to vehicle, whereas the positive control, an injection of semaglutide, resulted in almost 100 % taste aversion (**Fig 5J**). In addition, we measured water intake to understand if the CS powder containing sodium counterions affected the drinking behavior of the mice, something that has been observed with some metabolite salts^42^. Importantly, we observed no treatment-dependent differences in water intake (**Fig 5K-L**). This lack of hyperdipsia by CS treatment indicates that the sodium within the CS powder is unlikely to cause any obvious osmolarity-related problems *in vivo*, consistent with our behavioral examinations of CS treatment. These data suggest that the improved glucose tolerance observed in response to CS treatment is not driven by a sickness response induced e.g. by endotoxin-contamination, excess sodium, and/or viscosity of the commercial CS powder.

### Chondroitin sulfate improves glucose tolerance independently of effects on energy balance

Female mice fed a high-fat diet supplemented with CS^44^ and male mice fed a high-fat diet in combination with daily oral administration of fucosylated CS^45^ have been reported to weigh less than control-treated mice. Thus, we aimed to investigate if the improvement in glucose tolerance in DIO mice by subcutaneous CS treatment (**Fig 2-4**) was dependent on effects on energy balance regulation in DIO mice. We administered CS or a vehicle solution by daily subcutaneous injections for 14 days to DIO mice housed in metabolic cages (**Fig 6**). We found no treatment-dependent effects on body weight measurements (**Fig 6A-B**). However, DIO mice administered CS tended to consume fewer calories when compared to mice injected with a vehicle solution (**Fig 6C-D**). In line with the insignificant effects of subcutaneous CS compared to vehicle treatment on body weight loss in this study (**Fig 6A-B**), we found no differences in energy expenditure depending on treatment (**Fig 6E-F**). These data are consistent with placebo-controlled studies in humans showing no effects of oral CS on weight change from baseline, body composition, and resting energy expenditure in people with overweight and osteoarthritis compared to placebo^46,47^. Further, we found that mice treated subchronically with CS compared to vehicle tended to have decreased respiratory exchange ratios, especially during the dark phase (**Fig 6G-H**), indicating a slight increase in whole-body fat oxidation. These results suggest that the decreased glucose levels during glucose tolerance tests in CS compared to vehicle-treated DIO mice are not driven by increased glucose oxidation. Moreover, the lack of significant hypophagia and weight loss in response to CS treatment further underscore that the improved glucose tolerance seen in DIO mice is not driven by sickness.

**Figure 6.**
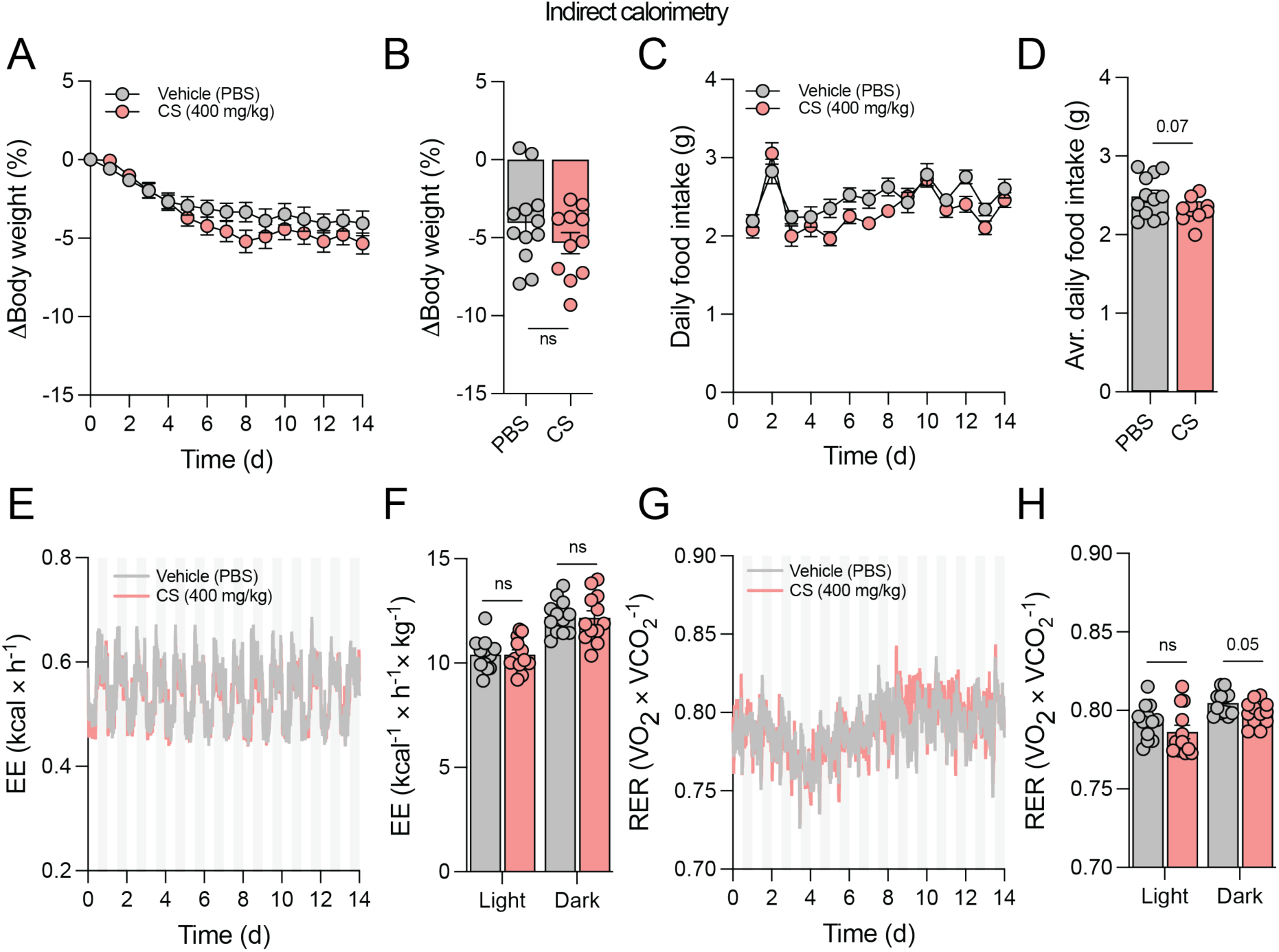
Subchronic subcutaneous chondroitin sulfate administration does not affect energy balance in diet-induced obese mice. **A-B)** Body weight change in percentage throughout the experiment (panel A) or at the end of the study (panel B). DIO mice (*n* = 12 per group) were subcutaneously injected with vehicle (grey) (PBS) or CS (red) daily for 14 days in metabolic cages (mouse study 9 in Materials and Methods). **C-D)** Daily food intake throughout the study (panel C) and as a daily average (panel D). **E-F)** Average energy expenditures (EE) throughout the study (panel E) or averages separated by light (left two bars) and dark (right two bars) phases (panel F). **G-H)** Average respiratory exchange ratios (RER) throughout the study (panel G) or averages separated by light (left two bars) and dark (right two bars) phases (panel H). Data presented as means +/-SEM and individual data points represent data from individual animals. Statistics in panels B, D, F, and H by unpaired t-test with Welch’s correction.

### Chondroitin sulfate decreases blood glucose levels in leptin-receptor deficient mice

To explore the glycemic effects of CS in other models of diabetes, we administered CS or vehicle subcutaneously to obese and severely hyperglycemic leptin-receptor deficient (*db^-^/db^-^*) mice and followed their blood glucose levels over two days (**Fig 7A**, **Fig S8A**). CS potently decreased blood glucose levels in diabetic and obese *db^-^/db^-^* mice over 24 h (**Fig 7A-B**, **Fig S8A**), and a similar effect was replicated in another cohort of *db^-^/db^-^* mice (**Fig S8B-C**). This acute effect was most pronounced at 4.5 h after the subcutaneous injection with around 50 % reduction in blood glucose levels in CS compared to vehicle-dosed *db^-^/db^-^* mice (**Fig 7A**). It is worthy to note that CS did not cause hypoglycemia but simply lowered blood glucose from ∼18 mM to ∼8 mM. This beneficial effect was not observed in obese leptin-deficient (*ob^-^/ob^-^*) mice (**Fig S8D-E**), suggesting that leptin is required for the effects of CS on blood sugar lowering. Additionally, the lack of effects in *ob^-^/ob^-^*mice again supports that the blood glucose lowering effects are not due to sickness from toxic contaminants^40,41^, viscosity, or an excess of co-administered sodium^42^.

**Figure 7.**
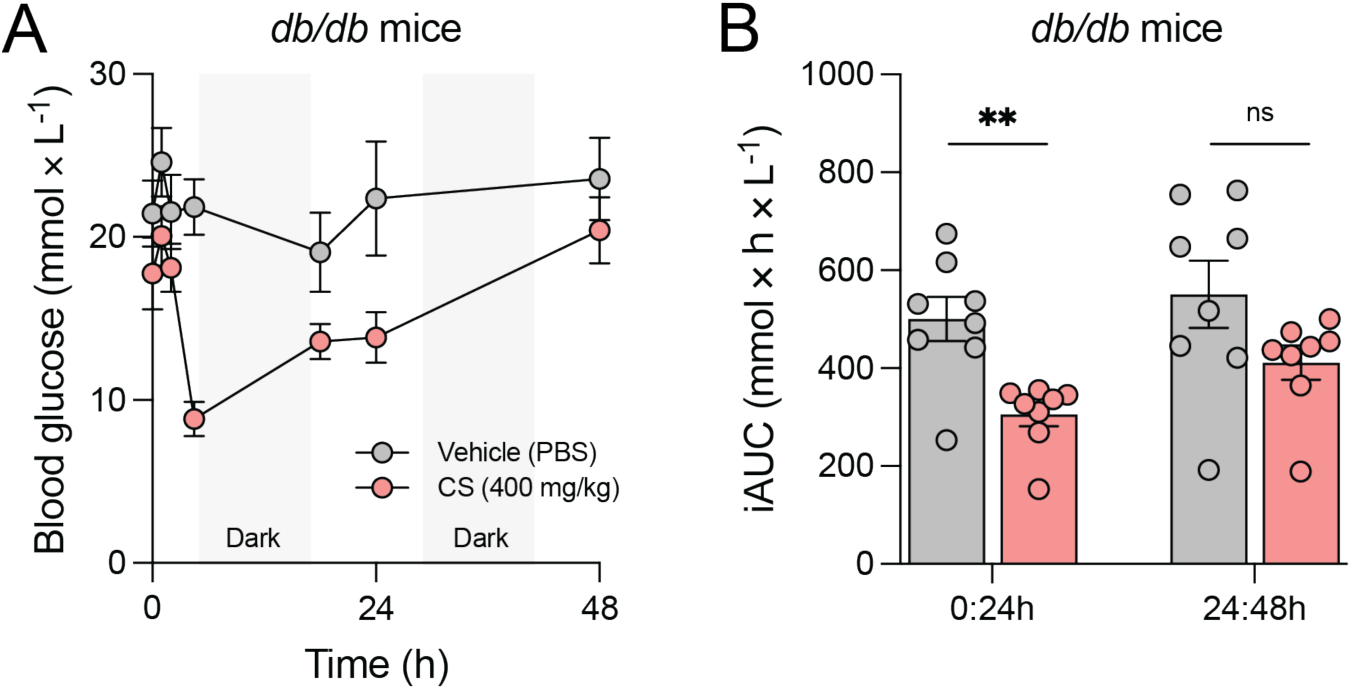
CS improves glycemic control in diabetic mice. **A-B)** Diabetic *db^-^/db^-^* mice (*n* = 8 per group) were subcutaneously injected with vehicle (grey) (PBS) or CS (red) and their blood glucose levels were measured throughout 48 h (panel A) and incremental areas under the curve (iAUC) were calculated (panel B) (mouse study 12 in Materials and Methods). Dark phases are indicated by light grey shaded areas (panel A). Statistics in panel B by unpaired t-tests with Welch’s correction. **, *p* < 0.01; ns, not significant.

This delayed reduction in blood glucose observed to occur later than 4 h after CS treatment in *db^-^/db^-^*mice (**Fig 7**) aligns with our previous observation that improved glucose tolerance depended on the timing of CS administration in DIO mice. In sum, these data demonstrate that subcutaneous administration of CS potently decreases basal blood glucose levels in diabetic *db^-^/db^-^* mice without causing hypoglycemia.

## Discussion

The pathophysiology of T2D is characterized by a series of metabolic defects that drive hyperglycemia. This includes insulin resistance and impaired insulin secretion but also low-grade inflammation and exacerbated lipolysis in adipose tissue. Research on the molecular mediators of these defects has largely focused on alterations in endocrine and intracellular signaling pathways, while less attention has been paid to the ECM and its potential role in the pathogenesis of T2D. Several lines of evidence suggest that the ECM is implicated in the development of insulin resistance and cardiometabolic disorders^14,48,49^. Studies in rodents and non-human primates show that impaired glycemic control is closely linked to remodeling of the ECM^50,51^. At the cellular level, ECM has also been reported to regulate insulin signaling in fat cells^52^. These findings suggest that maladaptive ECM remodeling is a factor that complements the classical ‘ominous octet’ of pathophysiological drivers of T2D. In line with this, we here identify robust links between genes related to the metabolism of one abundant ECM component, CS, and glycemic traits in humans. Moreover, we demonstrate that pharmacological doses of CS profoundly lower blood glucose in *db^-^/db^-^*mice and improve glucose tolerance and cardiac glucose clearance in diet-induced obese mice. This improved glycemic control is associated with an increased glucose-stimulated insulin secretion and improved insulin action and is not driven by adverse effects caused by e.g. endotoxemia. Together, these findings implicate the ECM in the pathogenesis of T2D and encourage further evaluation of matritherapies for managing hyperglycemia.

Our genetic analysis revealed significant associations between 12 CS-related human genes and both glycemic traits and T2D risk, providing new insights into the role of CS metabolism in these conditions. These genes are involved in various branches of CS metabolism, including CS biosynthesis (*GALNS*, *CSGALNACT1*, *CSGALNACT2, XYLT1*, *DSE*, *UST*, and *DSEL*), sulfation (*CHST10* and *CHST12*), and degradation (*HYAL4*, *GUSB*, and *ARSB*) of CS. While genetic studies have previously implicated ECM remodeling in the pathogenesis of type 2 diabetes and its cardiovascular comorbidities^53–58^, our results provide the first genetic links between CS genes and glycemic traits and T2D risk. Given that human genetic evidence is known to lower the risk of failure in drug development^28,59^, these findings provide impetus for further exploring CS biology as an anti-diabetic drug target.

Unlike in our studies, CS has previously been investigated for anti-diabetic effects in streptozotocin-induced diabetic rodents, mimicking ß cell loss of function or mass^25,26,38,60^, by administering CS orally^25,26,38,61^. For example, a single oral CS administration decreased blood glucose levels during glucose tolerance tests in both normal and strepto-zotocin-induced diabetic mice^38^. Additionally, streptozotocin-induced diabetic rats exhibited almost 50 % decreased basal blood glucose levels after 10 weeks of daily oral CS administration compared to control-treated streptozotocin rats^25^. Daily oral CS administered for 8 weeks in streptozotocin-induced diabetic rats decreased basal blood glucose levels by around 25 %^26^. Further, a single oral low molecular weight CS administration decreased blood glucose levels during oral sucrose and starch challenges, and daily CS administration decreased basal blood glucose levels after 3 weeks in a strain of albino mice compared to control-treated animals. While these previous studies provided insights into how orally administered CS regulates postprandial and basal blood glucose levels in streptozotocin-induced diabetic animals, our studies demonstrate that subcutaneous CS administration increases glucose-stimulated insulin secretion and glucose-induced glucose clearance in DIO mice.

Our work uncovers that subcutaneous CS administration to *db^-^/db^-^* mice leads to a dramatic decrease in blood glucose levels. The mechanism underlying this profound reduction in blood glucose in *db^-^/db^-^* mice is unclear, but it might be related to the decreased endothelial expression of *Csgalnact1* and the partly degraded endothelial glycocalyx previously observed in *db^-^/db^-^* mice compared to non-diabetic controls^33,62^. Moreover, disaccharide analysis of CS from the renal cortex of *db^-^/db^-^* mice have shown a reduction in 4-O-sulfated and 6-O-sulfated disaccharides and an increased content of unsulfated disaccharides^63^. Also, treatment with another sulfated polysaccharide, fucoidan, has been shown to reduce fasting plasma insulin levels and whole-body insulin resistance in *db^-^/db^-^* mice^64^.These effects were associated with a normalization of the degraded endothelial glycocalyx depth^62^, perhaps suggesting that sulfated ECM components, like fucoidan and CS, improves glycemic control in *db^-^/db^-^* by alleviating insulin resistance through a mechanism that might involve the density of endothelial glycocalyx.

Our findings indicate that the beneficial effect of subcutaneous CS on glycemic regulation is at least partly driven by improved insulin action. The proposed mechanisms for the reported anti-diabetic effects of orally administered CS include inhibition of α-amylase^38^ and α-glucosidase activities^61^, while also improvement of the action of insulin has been reported to be a potential mode of action for CS^65–68^. As α-amylase and α-glucosidase enzymes are primarily found in saliva, exocrine secretions from the pancreas, and on the microvilli of the gastrointestinal tract, the direct inhibition of these enzymes in the gastro-intestinal system is unlikely to account for the improved glucose tolerance in DIO mice administered CS by the subcutaneous route, as reported in our findings. Since oral CS supplementation has also been suggested to improve insulin action^65–68^, it is possible that oral and subcutaneous CS supplementations share some mechanisms while differing in others in improving blood glucose levels in DIO mice.

Given our observation of increased glucose-stimulated insulin secretion by CS, it was surprising that we found no evidence of increased glucose-stimulated glucose clearance by e.g. skeletal muscle. In contrast, we found that CS enhanced cardiac clearance of circulating glucose. This finding might point towards an attractive pharmacodynamic feature of chondroitin sulfate. Impaired metabolic flexibility in the heart (e.g. cardiac insulin resistance and impaired glucose oxidation) is a less appreciated aspect of the metabolic derangements seen in obesity and T2D. In line with this, enhancing glucose uptake and oxidation in the heart is linked to cardioprotective effects in the context of T2D^69^. Together, these observations suggest that pharmacological doses of CS exert anti-diabetic effects by restoring cardiac glucose uptake and metabolism. In this way, CS might complement the mechanisms of action for currently available T2D medications, illustrated by SGLT2 inhibitors that increase renal glucose excretion and metformin that primarily target the intestines, liver and skeletal muscle. However, we cannot exclude that CS also affects hepatic and urinary glucose output, as we did not investigate this in our experiments. In intraperitoneal glucose tolerance tests in mice, where insulin secretion is limited^70^, and considering that the liver is more sensitive to lower insulin concentrations compared to other peripheral tissues, inhibition of hepatic glucose output by CS may be a considerable factor in explaining the blood glucose-lowering effect of CS. The exact mechanism by which CS improves glucose tolerance in obese mice remains to be fully elucidated.

CS is a molecule with structural diversity in terms of its sulfation patterns and chain length^71^, two factors that are both known to determine the biological functions of this glycosaminoglycan^72^. In this study, we examined the effects of CS purified from shark cartilage, as it has been reported to improve postprandial blood glucose levels in streptozoto-cin-induced diabetic mice^38^. Moreover, shark cartilage is a common source of CS in nutritional supplements for humans^73–76^. However aside from shark-derived CS, CS compounds from other species, including pig^38^, cattle^26,38^, salamander^25^, and marine invertebrates^60,65–68^, amongst others, have been investigated for their anti-diabetic effects. These different sources of CS, combined with various isolation protocols and variations in purification and production strategies, contribute to the naturally occurring structural diversity of CS, stressing the complexity of studying CS^77–82^. Adding to this complexity are variations in both the administered doses and the routes of administration reported in the literature. Further, the quality of animal-derived CS isolations has been suggested to be poor and variable, where contaminants include immunogenic keratan sulfate and hyaluronic acid, raising interest in biotechnological, non-animal-derived CS^83^. Additionally, the heterogeneity of CS compounds in the treatment of osteoarthritis has been debated due to discordant endpoint efficacies reported^23,73^. In summary, inconsistencies in the sources, isolations, and purities of evaluated CS compounds might explain discrepant findings in the literature regarding efficacies, modes of action, and side effects.

In addition to its anti-diabetic effects, CS has been reported to have cardiometabolic benefits when administered exogenously for cardiovascular disease. CS administered in mice has been shown to protect against atherosclerosis and reduce adipose tissue mass^44,45,84–86^, while human studies have linked oral supplementation with CS to decreased serum cholesterol^87^, a reduced risk of acute myocardial infarction^87–91^ and ischemic stroke^92^, and lower cardiovascular mortality^87,89–91,93,94^. These findings illustrate that developing CS-based matritherapies has potential to not only improve glycemic control but also counteract the development of diabetic complications, thereby providing multi-modal treatments for improving the quality of life of people living with T2D. Additionally, advancements in the production of cell-based, specific, and pure protein-free GAGs or GAG-bearing proteoglycans^95,96^ might allow developments of inexpensive CS-based drugs and dietary supplements for treatment of T2D in low-income countries.

Lastly, CS from shark has a different sulfation pattern than human CS, but this should not discourage the field from investigating the “drugability” of CS biology. Both existing drugs and new molecular targets for the treatment of type 2 diabetes have previously emerged from studies of exotic animals, such as the venomous Gila monster^97^ and fish-hunting cone snails^98^.

In conclusion, our studies link CS biology to glycemic traits and T2D risk in humans and demonstrate that pharmacological doses of exogenous CS provide robust improvements in glycemic control without causing hypoglycemia or other adverse effects in mice. These anti-diabetic effects are likely driven by an enhanced cardiac glucose clearance, an improved insulin action, and an increase in insulin secretion that seem to be independent of the incretin system. These appealing pharmacodynamic properties, combined with intriguing human genetic evidence, highlight chondroitin sulfate as an attractive target for launching medicinal chemistry campaigns to develop novel anti-diabetic drugs that complement existing therapeutic options by acting upon an aspect of cell biology that is largely overlooked in the pathogenesis of type 2 diabetes.

## Author contributions

The project was initiated by J.L. and C.C. and further conceptualized together with V.B.I.J., A.W.B, and K.T-B.S. Human genetic analyses were conducted by M.J.R-L. and T.O.K. Mouse studies was conducted by V.B.I.J., A.W.B., C.S., J.L, K.SJ. and A.M.F. Tissue and plasma analyses were performed by V.B.I.J., K.SJ. and A.M.F. The manuscript was written by V.B.I.J. and J.L. All authors interpreted data and reviewed and approved the final manuscript.

## Declaration of interests

C.C. is a co-founder of Ousia Pharma, a biotech company developing therapeutics for treatment of obesity.

## Acknowledgments

V.B.I.J. was supported by a Peter and Emma Thomsen Foundation scholarship and by an NNF CBMR international PhD scholarship. J.L. was supported by the BRIDGE Trans-lational Excellence Programme (www.bridge.ku.dk) funded by the Novo Nordisk Foundation (grant number NNF20SA0064340). M.J.R-L was supported by a research grant from the Danish Diabetes and Endocrine Academy and the Danish Cardiovascular Academy, which are funded by the Novo Nordisk Foundation, grant numbers NNF22SA0079901 and NNF20SA0067242. R.L.M. was supported by the NovoNordisk Foundation grant NNF22OC0073736. T.O.K. was supported by the Novo Nordisk Foundation grant NNF22OC0074128. K.T.S is supported by a Novo Nordisk Foundation Hallas Møller Ascending Investigator grant (NNF0073793), the Carlsberg Foundation (CF21-0453) and a Sapere Aude Research Leader Grant from the Independent Research Fund Denmark (2066-00043B). The Novo Nordisk Foundation Center for Basic Metabolic Research is an independent Research Center, based at the University of Copenhagen, Denmark, and is partially funded by an unconditional donation from the Novo Nordisk Foundation (www.cbmr.ku.dk; grant numbers NNF18CC0034900 and NNF23SA0084103). The authors thank Dr. Jenny Brown for reading the manuscript and providing comments. We acknowledge the Rodent Metabolic Phenotyping Platform at the Novo Nordisk Foundation Center for Basic Metabolic Research (CBMR) for the technical expertise and support.

## Abbreviations

CS: Chondroitin sulfate
cis-eQTL: cis-expression quantitative trait locus
DIO: diet-induced obesity
ECM: extracellular matrix
GWAS: genome-wide association studies
MAF: ma-jor allele frequency
OTG: Open Targets Genetics
PIP: posterior inclusion probability
pLoF: predicted loss-of-function
cS2G: SNP-to-gene
T2D: type 2 diabetes
V2G: var-iant-to-gene

## Materials

### Human genetics

We employed both variant-level and gene-level approaches to identify genetic associations between 28 CS genes and T2D risk and glycemic traits, including fasting glucose levels, random glucose levels, and HbA_1c_ levels^99–102^. To examine whether genes encoding enzymes involved in chondroitin sulfate metabolism influence glycemic regulation in humans, we curated a panel of 28 genes implicated in CS metabolic pathways^30^ (**Fig. S1**, **Table S1**). This panel comprised 13 genes associated with CS biosynthesis, 9 with sulfation, and 6 with degradation.

### Variant-level analysis

We analyzed variant-level associations for 28 CS genes using two complementary integrative approaches. First, we identified common variants within ±1 Mb of each gene’s transcription start site and conducted fine-mapping to prioritize potential causal SNPs, which were subsequently mapped to their respective genes. Second, we explored the impact of rare coding variants in CS genes using data from the Genebass whole-exome sequencing database.

In the variant-level analysis for common variants (MAF > 1%), we could only use 27/28 genes due to CHST7 being present on chromosome X and not all glycemic GWAS having information on this chromosome. We first defined a region ±1 Mb from the transcription start site of each CS gene, mapped using biomaRT (v. 2.54.1) and tracklayer (v. 1.58.0). We then identified genome-wide significant (P < 5x10^-8^), independent (r^2^ < 0.01 and 1000 kb distance) lead genetic variants for T2D and glycemic traits in Europeans in these regions using the ieugwasr package (v. 0.1.5). To prioritize putative causal SNPs, we performed fine-mapping using CARMA^103^, extending the regions around the lead SNPs by ± 500 kb. To investigate whether these putative causal variants (posterior inclusion probability (PIP) > 0.1), are connected to the 28 CS-related genes, we mapped the variants to genes using combined SNP-to-gene (cS2G)^104^ and Open Target Genetics (OTG)^105^ approaches. The cS2G approach encompasses seven strategies: exon, promoter, eQTLGen blood *cis*-expression quantitative trait locus (eQTL), GTEx *cis*-eQTL, EpiMap enhancer-gene linking, ABC, and Cicero blood/basal. The OTG platform integrates distance to TSS, *cis*-eQTLs, protein QTLs, splicing QTLs, chromatin interactions, and *in silico* functional predictions. In the cS2G approach, we did not restrict the analysis to the best gene for each variant (cS2G after normalization > 0.5) since some of these genes are non-coding proteins, while others with cS2G after normalization < 0.5 are protein-coding. When testing the fine-mapped variants in OTG, we employed the R package otargen (v. 1.1.0)^106^ to retrieve the nearest gene information using the variantInfo() function, and used the genesForVariant() function to get the gene with the maximum V2G score for each variant. Finally, only lead variants or their proxies (r2 > 0.8) connected to the CS-related genes were reported. Proxies were found using the R package LDlinkR (v. 1.3.0).

We identified rare coding variants in CS genes by utilizing the Genebass whole-exome sequence database across 394,841 individuals from the UK Biobank^107^ (https://app.gene-bass.org/), which encompasses 8,074,858 coding variants tested across 4,529 phenotypes. In our single variant analysis, we obtained genome-wide significant (P < 5x10^-8^) rare-variant associations for each trait of interest. The single variant association results were downloaded from Genebass’s Hail library (gs://ukbb-exome-public/500k/results/results.mt) using Hail (https://hail.is/).

To further explore associations between the identified common variants and a wide range of glycemic phenotypes, we downloaded and curated the largest and latest GWAS in Europeans on glycemic traits, including random glucose^102^, insulin fold change^108^, pro-insulin^109^, 2-hour glucose, fasting glucose, fasting insulin and HbA_1c_^99^, and HOMA-B and HOMA-IR^110^. We aligned the effects to be positive in the GWAS where the common variant was found and tried to find the exact match or a proxy r2 > 0.8 in each glycemic GWAS. Proxies were found using the R package LDlinkR (v. 1.3.0). When possible, BMI-adjusted traits were used.

### Gene-Level Analysis

We evaluated gene-level associations for the 28 CS genes using two complementary approaches. First, we used Genebass to assess gene-based associations for coding variants. Second, we used MetaXcan to investigate the associations of genetically predicted mRNA expression levels of each CS gene with glycemic traits and T2D across 49 GTEx tissues^111^.

In our gene-level analysis for coding variants, we accessed gene-level association statistics for glycemic traits and T2D risk across three sets of variants (predicted loss-of-function, missense, synonymous) and three gene-based tests (Burden, SKAT, SKAT-O). We corrected these results for the number of genes tested (P < 0.05/28). In a similar fashion as in the single variant association analyses, the gene-based association results were downloaded from Genebass’s Hail library (gs://ukbb-exome-public/500k/results/re-sults.mt) using Hail (https://hail.is/).

In our gene-level analysis using non-coding variants, we identified CS genes for which genetically predicted mRNA expression is associated with glycemic traits or T2D risk. We employed MetaXcan to compute these gene-level association results across 49 human GTEx tissues^111^. We corrected the threshold of statistical significance for the number of genes and tissues tested (P < 0.05/(28*49)). The plots were created using gglot2 (v. 3.5.1). All analyses were conducted using R version 4.3.3.

### Animal studies

All animal studies were conducted in C57BL6/J male mice (Janvier Labs) unless otherwise stated. Upon arrival to the animal facility, mice were fed a chow diet (Altromin, 1310) or HFHS diet (58 kcal% fat, D12331i, Research Diets) from 8 weeks of age to maintain leanness or induce diet-induced obesity (DIO), respectively. For maintenance housing before onset of experimental procedure, chow-fed or DIO mice were housed in groups of 2-8 in closed, individually ventilated cages in a housing room with 12 h light-dark cycles (light phase: 06:00-18:00; dark phase: 18:00-06:00), at 22 °C ± 2 °C, and with ad libitum access to food and water, unless otherwise stated. For experiments, mice were double-housed unless otherwise specified. The animal experiments were performed in accordance with the Danish Animal Experimentation Inspectorate (animal license number: 2023-15-0201-01442).

### Study 1 – Effect of timing of subcutaneous chondroitin sulfate administration on glucose tolerance in diet-induced obese mice

Adult male DIO mice were administered either 400 mg/kg CS (concentration: 13.33 mg/mL CS in PBS, injection volume 30 µL/g body weight, Sigma Aldrich #9092-08-9) or PBS (injection volume 30 μL/g body weight) (*n* = 5 in each group). Injection volumes were distributed equally between two subcutaneous injections, one in each groin. Administration of CS was performed 2, 3, 4, 5, or 6 hours before intraperitoneal injection of 1.5 g/kg D-glucose, and administration of PBS was at 2 hours before intraperitoneal injection of D-glucose. At the first injection (6h before D-glucose injection), remnants of food in all cages were removed. Blood glucose levels were measured using blood from the tail tip and a handheld glucometer (Abbott) at timepoint 0 (just before intraperitoneal injection of D-glucose), 15, 30, 60, and 120 minutes after i.p injection of D-glucose.

### Study 2 – Effect of subcutaneous chondroitin sulfate administration on intraperitoneal glucose tolerance in diet-induced obese mice

Male 61-week-old DIO mice were injected subcutaneously with either 400 mg/kg CS (concentration: 10 mg/mL CS in PBS, injection volume 40 μL/g body weight, Sigma Aldrich #9092-08-9) or PBS (injection volume 40 μL/g body weight) (*n* = 10 in each group) 6 h before an intraperitoneal injection of 1.5 g/kg D-glucose. Injection volumes were distributed equally between two subcutaneous injections, one in each groin. At the time of sub-cutaneous injection, food remnants were removed. Viscosity controls included male 61-week-old mice injected subcutaneously with 8 g/kg D-mannitol (concentration: 200 mg/mL D-mannitol in PBS, injection volume 40 μL/g body weight) or 400 mg/kg dextran (concentration: 10 mg/mL in PBS, injection volume 40 μL/g body weight) (*n* = 5 per group). Blood glucose levels were measured using blood from the tail tip and a handheld glucometer (Abbott) at timepoint -360 (just before subcutaneous injection of CS/PBS), 0 (just before intraperitoneal injection of D-glucose), 15, 30, 60, and 120 min after intraperitoneal injection of D-glucose. Mice were randomized to the two groups and potential differences in blood glucose means and variances at timepoint -360 min between the groups were blocked. Subcutaneous injections were performed in the morning just after the first measurement of blood glucose (-360 min).

### Study 3 – Effect of subcutaneous chondroitin sulfate administration on glucose-stimulated insulin secretion in diet-induced obese mice

Male 63-week-old DIO mice were injected subcutaneous with either 400 mg/kg CS (concentration: 10 mg/mL CS in PBS, injection volume 40 μL/g body weight, Sigma Aldrich #9092-08-9) or PBS (injection volume 40 μL/g body weight) (*n* = 8 in each group). Injection volumes were distributed equally between two subcutaneous injections, one in each groin. CS and PBS were administered 6 h before an intraperitoneal injection of 1.5 g/kg D-glucose (Sigma, injection volume: 5 μL/g body weight, solvent: isotonic saline). Blood glucose levels were measured using blood from the tail tip and a handheld glucometer (Abbott) at timepoint -360 (just before subcutaneous injection of CS/PBS), 0 (just before intraperitoneal injection of D-glucose), 2, 10, and 30 min after intraperitoneal injection of D-glucose. Mice were randomized to the two groups and potential differences in blood glucose means and variances at timepoint -360 min between the groups were blocked. Subcutaneous injections were performed in the morning just after the first measurement of blood glucose (-360 min). At the same time points, blood was drawn from the tail tip for plasma insulin measurements. Blood samples were spun down at 2,500 *g* for 10 min at 4 °C. Plasma was isolated from the supernatant and kept at -80°C until plasma insulin levels were measured using an insulin ELISA kit (#90080, Crystal Chem) according to the manufacturer’s instructions.

### Study 4 – Acute effect of subcutaneous chondroitin sulfate administration on blood glucose levels and plasma insulin levels in diet-induced obese mice

Male 47-week-old DIO mice were injected subcutaneously with either 400 mg/kg CS (concentration: 10 mg/mL CS in PBS, injection volume 40 μL/g body weight, Sigma Aldrich #9092-08-9) or PBS (injection volume 40 μL/g body weight) (*n* = 8 in each group). Injection volumes were distributed equally between two subcutaneous injections, one in each groin. Blood glucose levels were measured using blood from the tail tip and a handheld glucometer (Abbott) at timepoint 0 (just before the subcutaneous injection of CS/PBS), and again 2, 4, and 6 h after the injections. At the same time points, blood was drawn from the tail tip for plasma insulin measurements. Blood samples were spun down at 2,500 *g* for 10 min at 4°C. Plasma was isolated from the supernatant and kept at -80°C until plasma insulin levels were measured using an insulin ELISA kit (#90080, Crystal Chem) according to the manufacturer’s instructions. Mice were randomized to the two groups and potential differences in blood glucose means and variances at timepoint 0 h between the groups were blocked. Subcutaneous injections were performed in the morning just after the first measurement of blood glucose (0 h).

### Study 5 – Effect of subcutaneous chondroitin sulfate administration on insulin tolerance in diet-induced obese mice

Adult male DIO mice were injected subcutaneously with either 400 mg/kg CS (concentration: 13.33 mg/mL CS in PBS, injection volume 30 µL/g body weight, Sigma Aldrich #9092-08-9) or PBS (injection volume 30 μL/g body weight) (n = 5 in each group). Injection volumes were distributed equally between two subcutaneous injections, one in each groin. Administration of CS or PBS was performed 4 hours before intraperitoneal injection of insulin (Novo Nordisk, 0.75 U/kg body weight, 5 µL/g body weight, solvent: isotonic saline). Remnants of food were removed from cages at the time of the injection. Blood glucose levels were measured using blood from the tail tip and a handheld glucometer (Abbott) at timepoint 0 (just before intraperitoneal injection of insulin), 15, 30, 60, and 120 minutes after i.p injection of insulin.

### Study 6 – Effect of subcutaneous chondroitin sulfate administration on tissue-specific glucose-induced glucose clearance in diet-induced obese mice

Male 65-week-old DIO mice were injected subcutaneously with either 400 mg/kg CS (concentration: 10 mg/mL CS in PBS, injection volume 40 μL/g body weight, Sigma Aldrich #9092-08-9) or PBS (injection volume 40 μL/g body weight) (n = 10 in each group). Injection volumes were distributed equally between two subcutaneous injections, one in each groin. CS and PBS was administered 6 h before an intraperitoneal injection of 1.5 g/kg D-glucose (Sigma, 10 μL/g body weight, solvent: isotonic saline) and 60 μCi/ml ^3^H-la-beled-2-deoxy-glucose (#MT911, Hartmann Analytic GMBH). Blood glucose levels were measured using blood from the tail tip and a handheld glucometer (Abbott) at timepoint - 360 (just before subcutaneous injection of CS/PBS), 0 (just before intraperitoneal injection of 2-deoxy-glucose), 10, 20, 30, and 40 min after intraperitoneal injection of 2-deoxy-glucose. Mice were randomized to the two groups and potential differences in blood glucose means and variances at timepoint -360 min between the groups were blocked. Sub-cutaneous injections were performed in the morning just after the first measurement of blood glucose (-360 min). At the same time points, for analysis of ^3^H-2-DG clearance in indicated tissues, blood ^3^H activity was measured at 10, 20, and 40 min in 5 μl of blood by scintillation counting (Packard TriCarb 2900TR, Perkin-Elmer) and systemic ^3^H-2-DG exposure estimated by the trapezoidal method. At time point 40 min, mice were sacrificed by decapitation, trunk blood was collected in EDTA-coated tubes, and the following tissues were harvested in this order: gastrocnemius, quadriceps, heart, eWAT, iWAT, BAT, liver, and whole brain. Tissues were snap-frozen in liquid N_2_ and stored at -20 °C until measurements of radioactivity were performed. Trunk blood samples were spun down at 2,500 *g* for 10 min at 4°C. Plasma was isolated from the supernatant and kept at -80°C. To estimate the accumulation of ^3^H-2-DG-6-phosphate (^3^H-2-DG-6-P), a 15-50 mg sample of each tissue was used for the precipitation method^112^. Glucose clearance was calculated by dividing tissue ^3^H-2-DG-6-P counts by systemic ^3^H-2-DG exposure^113^. Western-blotting analyses of heart lysates were performed as described previously^114^. In brief, 30 mg of heart tissue was homogenized in ice-cold homogenization buffer, rotated end-over-end for 1 h, and lysates obtained after centrifugation at 12,000 g at 4 °C. Lysates were subjected to protein determination, diluted to the same protein concentration, and subjected to SDS-PAGE and immunoblotting. The primary antibodies were from Cell Signalling; Akt S473 (#9271, AB_329825), Akt T308 (#9275, AB_329828), Akt2 (#3063, AB_2225186), TBC1D4 T642 (#8881, AB_2651042), except TBC1D4 that were from Abcam (#ab189890, AB_2818964). Bands were visualized using a Bio-Rad ChemicDoc MP Imaging System (Bio-Rad, US).

### Study 7 – Effect of subcutaneous chondroitin sulfate administration on behavior of lean mice in an open field

Male 22-week-old lean mice fed a chow diet were single-housed in open cages and acclimatized to the procedure room one week before the initiation of the experiment. Mice were randomized to the two groups and potential differences in bodyweight means and variances between the groups were blocked. In the morning on the day of the experiment, mice were administered either 400 mg/kg CS (concentration: 10 mg/mL CS in PBS, injection volume 40 μL/g body weight, Sigma Aldrich #9092-08-9) or PBS (injection volume 40 μL/g body weight) (*n* = 6 in each group) by subcutaneous injection before the recording of their movement via ceiling-mounted Logitech C920 Pro cameras (1080 × 1080 pixels, 30 frames per second, Logitech software). Injection volumes were distributed equally between two subcutaneous injections, one in each groin. Their movements were recorded for 20 min after mice were placed in the middle of the arenas just after subcutaneous injections. Treatments were equally distributed in each run and across all chambers. Noldus EthoVision XT software (Noldus) was used to quantify locomotor activity based on their movement tracing algorithms.

### Study 8 - Effect of subcutaneous chondroitin sulfate administration on voluntary wheel running in lean mice

Male 28-week-old lean mice fed ad libitum chow diet and tap water were single-housed and acclimatized to open cages equipped with running wheels (23 cm in diameter, Techniplast activity cage, Techniplast) for 1 week. The amount of bedding was reduced to avoid any blocking of the running wheels. The running distance and time of mice were measured for 48 h before and after administration of either a subcutaneous injection of 400 mg/kg CS (concentration: 10 mg/mL CS in PBS, injection volume 40 μL/g body weight, Sigma Aldrich #9092-08-9) or PBS (injection volume 40 μL/g body weight) (*n* = 5 in each group). Injection volumes were distributed equally between two subcutaneous injections, one in each groin. Mice were randomized to the two groups and differences in means and variances of running distance and time between the groups were blocked. Running distance and time were measured by an odometer (Sigma Pure 1 Topline 2016, Sigma).

### Study 9 – Effect of sub-chronic subcutaneous chondroitin sulfate administration on energy balance and behavioral parameters in diet-induced obese mice

Male 39-week-old DIO mice were single-housed in metabolic cages (indirect calorimetry, Promethion Core^®^, Sable Systems). Mice acclimatized to the systems for 72 h before the study start and were randomized to either of the two groups: once-daily subcutaneous injections for 14 days of 400 mg/kg CS (concentration: 10 mg/mL CS in PBS, injection volume 40 μL/g body weight, Sigma Aldrich #9092-08-9) or PBS (injection volume 40 μL/g body weight) (*n* = 12 per group). Injection volumes were distributed equally between two subcutaneous injections, one in each groin. The two groups were equally distributed in both systems and within the system to the Promethion Core’s combined gas analyzers and flow regulation modules mounted to the environmental cabinets. Mice were injected daily between 4-6 PM, together with measurements of food intake and body weight. To minimize spillage of food, old pellets two new HFHS food pellets were provided each. Fresh tap water was provided once a week. Locomotor activity was measured by the Promethion beam break monitor. MacroInterpreter (Sable Systems) was used to output data on O_2_ consumption, CO_2_ production, respiratory exchange ratio, water intake, and locomotion at a resolution of every 5 min. Data from MacroInterpreter was imported into R, and relative time was transformed, such that time 0 was the beginning of the light phase on the first day of injection. Data was presented either as continuously smoothened over 20-minute intervals during the 14 days or averages over relative periods, reflecting dark and light phases.

### Study 10 – Effect of subcutaneous chondroitin sulfate administration on intraperitoneal body temperature

Male 22-week-old lean mice (*n* = 16) were single-housed and kept on ad libitum chow diet (SAFE D30) and tap water. One week later, mice had sterile RFID temperature transponders (UCT-2112 microchips, Unified Information Devices) surgically implanted for automatic measurements of core body temperature using the UID Mouse Matrix system (Unified Information Devices). Mice were anesthetized with isoflurane in oxygen. After induction of anesthesia, mice were placed on a heating pad kept at 37C throughout the entire surgical procedure. Incision sites on the abdomen were prepared by shaving and disinfection (70% ethanol following by 0.5% Chlorhexidine in 80% ethanol). Following this, mice received subcutaneous injection of Carprofen (5 mg/kg, Norodyl Vet, Scanvet) together with a combination of Lidocaine (7 mg/kg, Xylocain, AstraZeneca) and Bupivacaine (7 mg/kg, Marcaine, Orifarm) at the site of incision for local anesthesia. Next, mice were moved to the surgical field and covered with sterile drapes. Under aseptic conditions, an approximately 5 mm midline central abdominal incision was made followed by insertion of the temperature transponder into the abdominal cavity. Lastly, the incision was rinsed with sterile isotonic saline followed by closure of the skin with 7 mm wound clips (Reflex Autoclips, AgnThos). After surgery, all single-housed mice were placed in clean cages with access to a 37C heating pad for the subsequent 24 hours. Moreover, for two days following the surgery, mice were monitored to ensure proper wound healing and additional analgesia were provided (5 mg/kg carprofen). The wound clips were removed around 1 week after the surgery. At an age of 26 weeks, mice were treated with either 400 mg/kg CS (n=8, solvent: PBS, Sigma, #C4384, injection volume: 40 μL/g body weight administered as two equally sized subcutaneous injections in the inguinal area, concentration: 10 mg/mL) or PBS (n=8, injection volume: 40 μL/g body weight). Injection volumes were distributed equally between two subcutaneous injections, one in each groin. Injections were made in the afternoon around 5PM. Body temperature data was collected automatically at a time resolution of 5 min both before and after the experimental injections.

### Study 11 – Conditioned taste aversion of subcutaneous chondroitin sulfate in lean mice

Male 34-week-old lean mice fed a chow diet were habituated to ventilated cabinet scantainers (Scanbur) in single-housed open cages with normal daylight cycles at 22 ± 2 °C for 7 days before the experiment. The sunrise and sunset times were set to 20 min, the daylight level was 50 lux, and the relative humidity was 55 ± 20 % with 50 ± 10 air changes per hour. During the habituation period, mice were acclimatized to the presence of two water bottles in their home cage. Drinking bottles with metal nipples (Akronom, 84-ACCP2511) having smaller diameters of the opening compared to regular drinking were used to minimize leakage of liquid from the bottles. These drinking bottles are specifically designed for studies necessitating control of water consumption. Before the conditioning day, mice were water-deprived for 16 h (afternoon until the next morning) to ensure liquid intake during conditioning. On the conditioning day, mice were exposed to 0.1 % saccharine solution in both of their drinking bottles for 4 h in combination with either a subcutaneous injection of 400 mg/kg CS (concentration: 10 mg/mL CS in PBS, injection volume 40 μL/g body weight), isovolumic PBS, or 10 nmol/kg semaglutide (MedChemExpress, Cat. No.: HY-114118, vehicle: 8 mM PBS and 240 mM propylene glycol). For CS and PBS, injection volumes were distributed equally between two subcutaneous injections, one in each groin. For semaglutide, mice were treated with a single injection at 5 μL/g body weight in one groin. Subsequently, mice had access to normal water for 36 h. Mice were randomized to either of the three groups (*n* = 10 per group) based on the baseline water intake and body weight before conditioning. On the preference test day, mice had access to one bottle of 0.1 % saccharine and one with normal water. Side preference was blocked by alternating the position of saccharin between the right and left sides of the cages. Saccharin and water intake were measured throughout 24 h.

### Study 12 – Effect of subcutaneous chondroitin sulfate administration on blood glucose in *db^-^/db^-^*mice

Male *db^-^/db^-^* mice (strain BKS(D)-Lepr db/dbJOrlRj from Janvier Labs) fed chow diet (SAFE D30) were injected subcutaneously with either 400 mg/kg body weight chondroitin sulfate (concentration: 13.33 mg/mL PBS, injection volume: 30 µL per g body weight, Sigma Aldrich #C4384-5G) or PBS (injection volume: 30 µL per g body weight). Injection volumes were distributed equally between two subcutaneous injections, one in each groin. Blood levels of glucose were measured by applying a drop of whole blood from the tail tip onto a glucose strip inserted into a handheld glucometer (Abbott). The average of two blood glucose measurements was used as data points. A blood glucose value of 35 mmol/L was used in the few cases where blood glucose was above the range measurable by the glucometer. In this study, 16 *db^-^/db^-^* mice at 7 weeks of age weighing between 37 g and 43 g were divided into two groups with similar average body weight (40 g) and blood glucose (19 mmol/L). Eight mice were injected with PBS and seven of those were severely diabetic (baseline blood glucose values of 25.0, 17.9, 23.9, 20.2, 11.2, 22.8, 23.4, and 8.2 mmol/L the day before the experiment). The other eight mice were treated with CS and they were all severely diabetic (blood glucose levels of 24.3, 26.7, 21.0, 13.9, 13.7, 19.2, 14.0, and 18.8 mmol/L the day before the experiment). Injections were made in the afternoon around 5 h before the onset of the dark phase. Blood glucose was measured before injections (0 h) and 1, 2, 4.5, 18, 24, and 48 h after injections.

### Study 13 – Effect of subcutaneous chondroitin sulfate administration on blood glucose in *db^-^/db^-^*mice

Male *db^-^/db^-^* mice (strain BKS(D)-Lepr db/dbJOrlRj from Janvier Labs) fed chow diet (SAFE D30) were injected subcutaneously with either 400 mg/kg chondroitin sulfate (concentration: 13.33 mg/mL PBS, injection volume: 30 µL per g body weight, Sigma Aldrich #C4384-5G) or PBS (injection volume: 30 µL per g body weight). Injection volumes were distributed equally between two subcutaneous injections, one in each groin. Blood glucose was measured using blood from the tail tip and a handheld glucometer (Abbott). The average of two blood glucose measurements was used as data points. A blood glucose value of 35 mmol/L was used in the few cases where blood glucose was above the range measured by the glucometer. In this pilot study, 9 *db^-^/db^-^* mice at 10 weeks of age weighing between 33 g and 49 g were used. Four mice were injected with PBS, and they were all severely diabetic (baseline blood glucose values of 24.8, 24.3, 18.1, and 27.3 mmol/L). Five mice were injected with CS and four of those were severely diabetic (baseline blood glucose levels of 16.7, 24.9, 29.8, 25.6, and 6.6 mmol/L). Injections were made in the afternoon around 3 h before the onset of the dark phase. Blood glucose was measured before injections (0 h) and 1, 2, 6, 17, 24, and 48 h after injections.

### Study 14 – Effect of subcutaneous chondroitin sulfate administration on blood glucose in *ob/ob* mice

Twenty obese leptin-deficient ob/ob mice were kept on ad libitum chow diet (SAFE D30) and tap water following arrival to the animal facility and housed at 22C (plus/minus 2 grader) and 12-h light-dark phases (light phase: 6am-6pm, dark phase: 6am-6pm). The mice were single-housed and split into two groups with similar average body weight (around 56 g) and mean resting blood glucose (around 7.8 mmol/L) in the morning on the day of the injections. Next, mice were treated with either 400 mg/kg CS (n=10, solvent: PBS, Sigma, Sigma Aldrich #9092-08-9, injection volume: 30 uL/ g body weight, concentration: 13.33 mg/mL) or PBS (n=10, injection volume: 30 uL/ g body weight). Injection volumes were distributed equally between two subcutaneous injections, one in each groin. Blood levels of glucose were measured by applying a drop of whole blood from the tail tip onto a glucose strip inserted into a handheld glucometer (Abbott). Following baseline blood sampling (0 h), injections were made in the afternoon around 1pm and blood glucose were measured again 1, 2, 4, 20, 24, and 48 h after injections.

### Study 15 – Effect of subcutaneous chondroitin sulfate on intraperitoneal glucose tolerance in diet-induced obese mice when administered 30 min or 3 h before glucose

Adult male DIO mice were administered either 400 mg/kg CS (concentration: 13.33 mg/mL CS in PBS, injection volume 30 µL/g body weight, Sigma Aldrich #9092-08-9) or PBS (injection volume 30 μL/g body weight) (*n* = 5-6 in each group). Injection volumes were distributed equally between two subcutaneous injections, one in each groin. Administration was performed at 3 hours and 30 minutes before intraperitoneal injection of 1.5 g/kg D-glucose. All animals were injected at both time points, receiving CS at either 3 hours or 30 minutes before D-glucose administration and PBS for the remaining time point. The PBS group was administered PBS at both time points. All remnants of food were removed from the cages 6 hours before intraperitoneal injection of D-glucose. Blood glucose levels were measured using blood from the tail tip and a handheld glucometer (Abbott) at timepoint 0 (just before intraperitoneal injection of D-glucose), 15, 30, 60, and 120 minutes after i.p injection of D-glucose.

### Study 16 – Effect of subcutaneous chondroitin sulfate administration the day prior to a glucose challenge on intraperitoneal glucose tolerance in diet-induced obese mice

Adult male DIO mice (*n* = 7-8 per group) were injected subcutaneously with 400 mg/kg CS (concentration: 13.33 mg/mL CS in PBS, injection volume 30 µL/g body weight, Sigma Aldrich #9092-08-9) or PBS (injection volume 30 μL/g body weight) (*n* = 7-8 in each group). Injection volumes were distributed equally between two subcutaneous injections; one in each groin. CS and PBS was administered 21 hours before an intraperitoneal injection of 1.5 g/kg body weight D-glucose. In the morning of the intraperitoneal glucose injection, i.e. 6 hours before the glucose injection, remnants of food in all cages were removed. Blood glucose levels were measured using blood from the tail tip and a handheld glucometer (Abbott) at timepoint 0 (just before intraperitoneal injection of D-glucose), 15, 30, 60, and 120 minutes after i.p injection of D-glucose.

### Study 17 – Effect of oral chondroitin sulfate administration on intraperitoneal glucose tolerance in diet-induced obese mice

Male 56-week-old DIO mice were treated by oral gavage with either 400 mg/kg CS (concentration: 80 mg/mL CS in PBS, gavage volume 5 μL/g body weight, Sigma Aldrich #9092-08-9) or PBS (gavage volume 5 μL/g body weight) (*n* = 8 in each group) by oral gavage 6 h before an intraperitoneal injection of 1.5 g/kg D-glucose (sigma, solvent: isotonic saline). At the time of subcutaneous injection, food remnants were removed. Blood glucose levels were measured using blood from the tail tip and a handheld glucometer (Abbott) at timepoint -360 (just before subcutaneous injection of CS/PBS), 0 (just before intraperitoneal injection of D-glucose), 15, 30, 60, and 120 min after intraperitoneal injection of D-glucose. Mice were randomized to the two groups and potential differences in blood glucose means and variances at timepoint -360 min between the groups were blocked. Oral gavage was performed in the morning just after the first measurement of blood glucose (-360 min).

### Study 18 – Effect of intracerebroventricular chondroitin sulfate administration on intraperitoneal glucose tolerance in lean mice

Adult male mice on standard chow diet were single housed in open cages. Mice were anaesthetized by isoflurane, fixed in a stereotaxic frame (Kopf Instruments), and injected subcutaneously with local anesthesia (Lidocaine, Accord Healthcare) at the incision site of the skull. Next, the skin above the skull was incised and a hole was drilled into the skull and a guide cannula (26GA, PlasticOne, C2315GS-4/SPC) was inserted into the lateral ventricle using the following stereotaxic coordinates: −0.3 mm posterior to and ±1.0 mm lateral to bregma. The guide cannula was held in place by ultraviolet-cured cement (G-bond and G-aenial Universal Flo, GC). To keep the cannula closed between infusions, a dummy cannula (PlasticsOne, C315DCS-4/Spc, 2.5 mm) was inserted. Mice were treated with 5 mg/kg body weight carpofen (Pfizer) on the day of the surgery and for 3 days postoperatively. Mice recovered for a minimum of 7 days. After recovery, correct cannula placement was tested by observing water drinking responsiveness to intracerebroventricular infusion of 1 µL human angiotensin II (Sigma-Aldrich, A9525) at a concentration of 24 µM in artificial cerebrospinal fluid (distilled water with 125 mM NaCl, 2.5 mM KCl, 2.6 mM Na-HCO3, 1.25 mM NaH2PO4·2H2O, 25 mM D-glucose monohydrate, 1 mM MgCl2, 2 mM CaCl2). Mice that did not drink water within 15 min of infusion were excluded from the study. At the day of the study, the mice were infused intracerebroventricularly with either CS (10 µg/µL, infusion volume 8 µL) or PBS (infusion volume 8 µL) 4 hours prior to intraperitoneal injection of 1.5g/kg D-glucose injection. Total infusion volume was divided in two and infused at 2 µL/min with 1 min pause in between infusions. Before the infusion, the dummy cannula was carefully removed, and infusion was done using a Hamilton syringe mounted an automated syringe pump (Harvard Apparatus) and a 33-gauge injector (PlasticsOne, C315IS-4/SPC, 2.5 mm). To ensure complete infusions and avoid backflow, the injector was kept in the guide cannula for 30–60 s after infusion stop. Blood glucose levels were measured using blood from the tail tip and a handheld glucometer (Abbott) at timepoint 0 (just before the intraperitoneal injection of D-glucose), and again 15, 30, 60, and 120 minutes after intraperitoneal injection of D-glucose.

### Study 19 – Effect of subcutaneous chondroitin sulfate administration on intra-peritoneal glucose tolerance in lean mice

Male 21-week-old lean mice fed a chow diet were injected subcutaneous with either 400 mg/kg CS (concentration: 10 mg/mL CS in PBS, injection volume 40 μL/g body weight, Sigma Aldrich #9092-08-9) or PBS (injection volume 40 μL/g body weight) (*n* = 12 in each group). Injection volumes were distributed equally between two subcutaneous injections, one in each groin. CS and PBS were administered 6 h (360 min) 6 h before an intraper-itoneal injection of 2 g/kg D-glucose (sigma, solvent: isotonic saline). Blood glucose levels were measured using blood from the tail tip and a handheld glucometer (Abbott) at timepoint -360 min (just before subcutaneous injection of CS/PBS), 0 min (just before intraperitoneal injection of D-glucose), and again 15, 30, 60, and 120 min after the intra-peritoneal injection of D-glucose. Mice were randomized to the two groups and potential differences in blood glucose means and variances at timepoint -360 min between the groups were blocked. Subcutaneous injections of CS and PBS were performed at 7-8 AM in the morning, just after the first measurement of blood glucose (-360 min).

### Study 20 – Acute effect of subcutaneous chondroitin sulfate administration on blood glucose levels and plasma insulin levels in lean mice

Male 22-week-old lean mice fed a chow diet were injected subcutaneously with either 400 mg/kg CS (concentration: 10 mg/mL CS in PBS, injection volume 40 μL/g body weight, Sigma Aldrich #9092-08-9) or PBS (injection volume 40 μL/g body weight) (*n* = 8 in each group). Injection volumes were distributed equally between two subcutaneous injections, one in each groin. Blood glucose levels were measured using blood from the tail tip and a handheld glucometer (Abbott) at timepoint 0 (just before the subcutaneous injections of CS/PBS), and again 2, 4, and 6 h after the injections. At the same time points, blood was drawn from the tail tip for plasma insulin measurements. Blood samples were spun down at 2,500 *g* for 10 min at 4°C. Blood plasma was isolated from the supernatant and kept at -80 °C until plasma insulin levels were measured using an insulin ELISA kit (#90080, Crystal Chem) according to the manufacturer’s instructions. Mice were randomized to the two groups and potential differences in blood glucose means and variances at timepoint 0 h between the groups were blocked. Subcutaneous injections were performed in the morning just after the first measurement of blood glucose (0 h).

### Study 21 – Effect of two daily subcutaneous chondroitin sulfate administration on energy balance in diet-induced obese mice

Male 39-week-old DIO mice were single-housed in metabolic cages (indirect calorimetry, Promethion Core^®^, Sable Systems). Mice acclimatized to the systems for 72 h before the study started and were randomized to either of the two groups: once-daily subcutaneous injections of 400 mg/kg CS (concentration: 10 mg/mL CS in PBS, injection volume 40 μL/g body weight, Sigma Aldrich #9092-08-9) or PBS (injection volume 40 μL/g body weight) (*n* = 8 per group) for 2 days. Injection volumes were distributed equally between two subcutaneous injections, one in each groin. The two groups were equally distributed in both systems and within the system to the Promethion Core’s combined gas analyzers and flow regulation modules mounted to the environmental cabinets. CS and PBS were administered between 03.00-05.00 PM where body weight was also measured. Assessment of locomotor activity was enabled by the Promethion beam break activity monitor. MacroInterpreter (Sable Systems) was used to output data on O_2_ consumption, CO_2_ production, respiratory exchange ratio, water intake, and locomotion at a resolution of every 5 min. Data from MacroInterpreter was imported into R, and relative time was transformed, such that time 0 was the beginning of the light phase on the first day of injection. Data was presented as averages over time during the treatment period.

**Figure S1.**
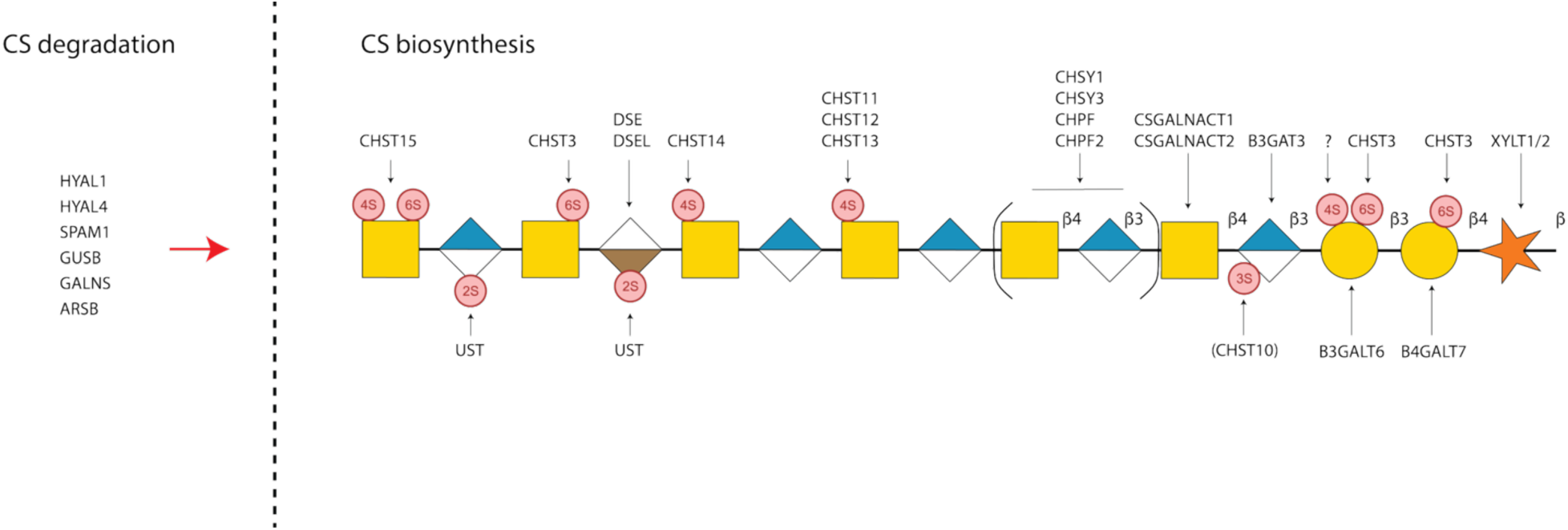
Schematic representation of the enzymatic steps involved in chondroitin sulfate (CS) degradation, biosyn-thesis, and sulfation. The enzyme products of the 28 CS-related genes analyzed in the present genetic screen are shown at the corresponding steps in the pathway. Yellow square: N-acetylgalactosamine. White and blue square: glu-curonic acid. White and brown square: iduronic acid. Yellow circle: galactose. Orange star: xylose. Small red circle: sulfation position indicated by alpha carbon atom numbering. Black lines: covalent bonds between sugar units. A com-plete list of all genes, their names, and annotated functions is provided in Supplementary Table 1. Related to main figure 1.

**Figure S2.**
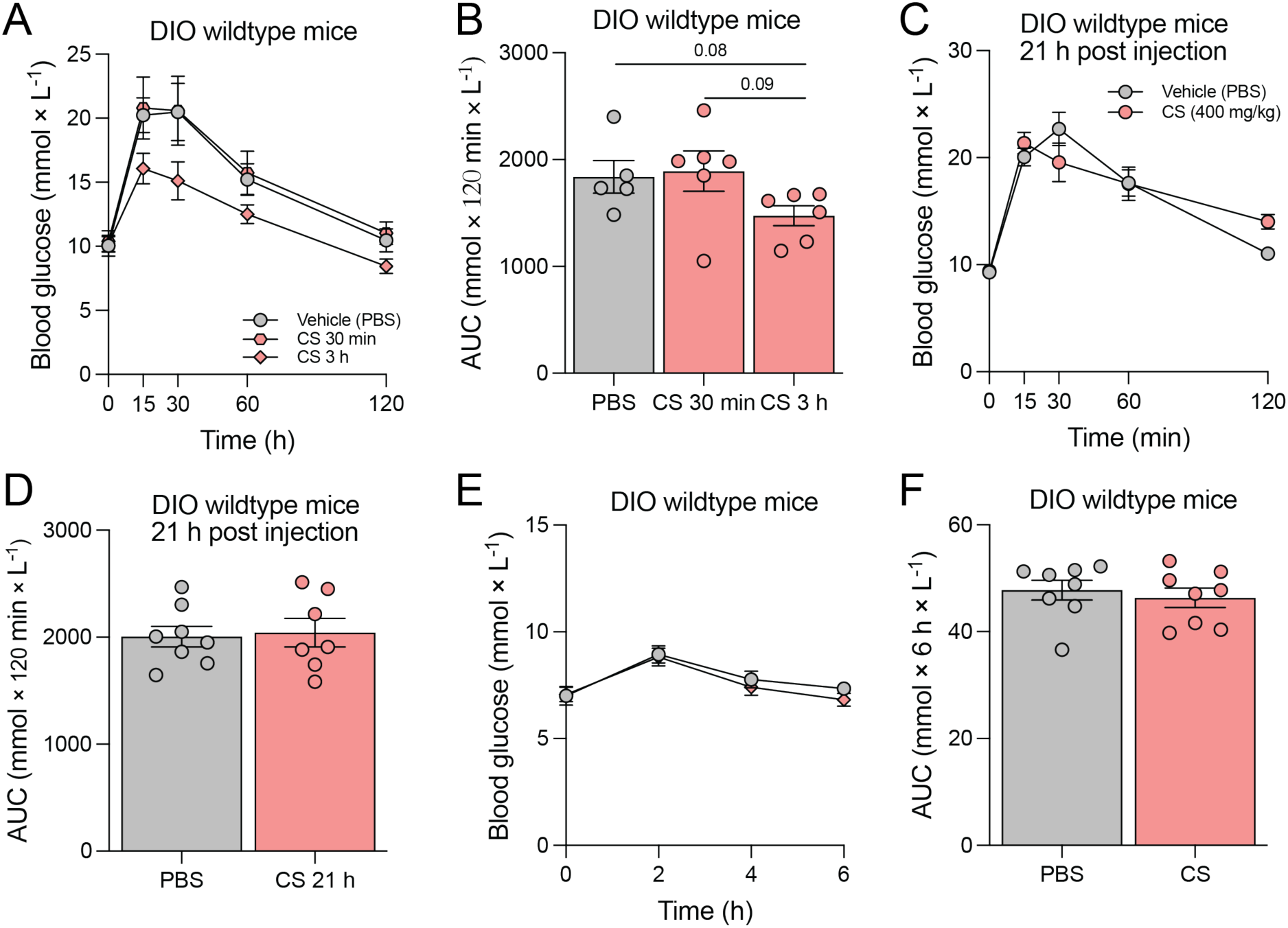
**A-B)** Glucose tolerance tests in diet-induced obese (DIO) mice with time dependency of CS dosing. DIO were subcutaneous injected with vehicle 3 h (grey) or CS either 30 min or 3 h (red hexagons and rhombi, respectively, and as indicated) before an intraperitoneal injection of glucose, and their blood glucose levels were measured (panel F) and the AUCs calculated (panel G) (*n* = 5-6 per group) (mouse study 15 in Materials and Methods). **C-D)** DIO mice (*n* = 7-8 per group) were injected subcutaneous with vehicle (grey) or CS (red) 21 h before an intraperitoneal injection of glucose, and their blood glucose levels were measured (panel H) and the AUCs calculated (panel I) (mouse study 16 in Materials and Methods). **E-F)** Acute compound tolerance test of CS vs. vehicle on blood glucose levels in DIO mice (panel J) and the measured AUCs (panel K) (*n* = 8 per group) (mouse study 4 in Materials and Methods). Statistics by one-way ANOVA in panel G. Related to main figure 2.

**Figure S3.**
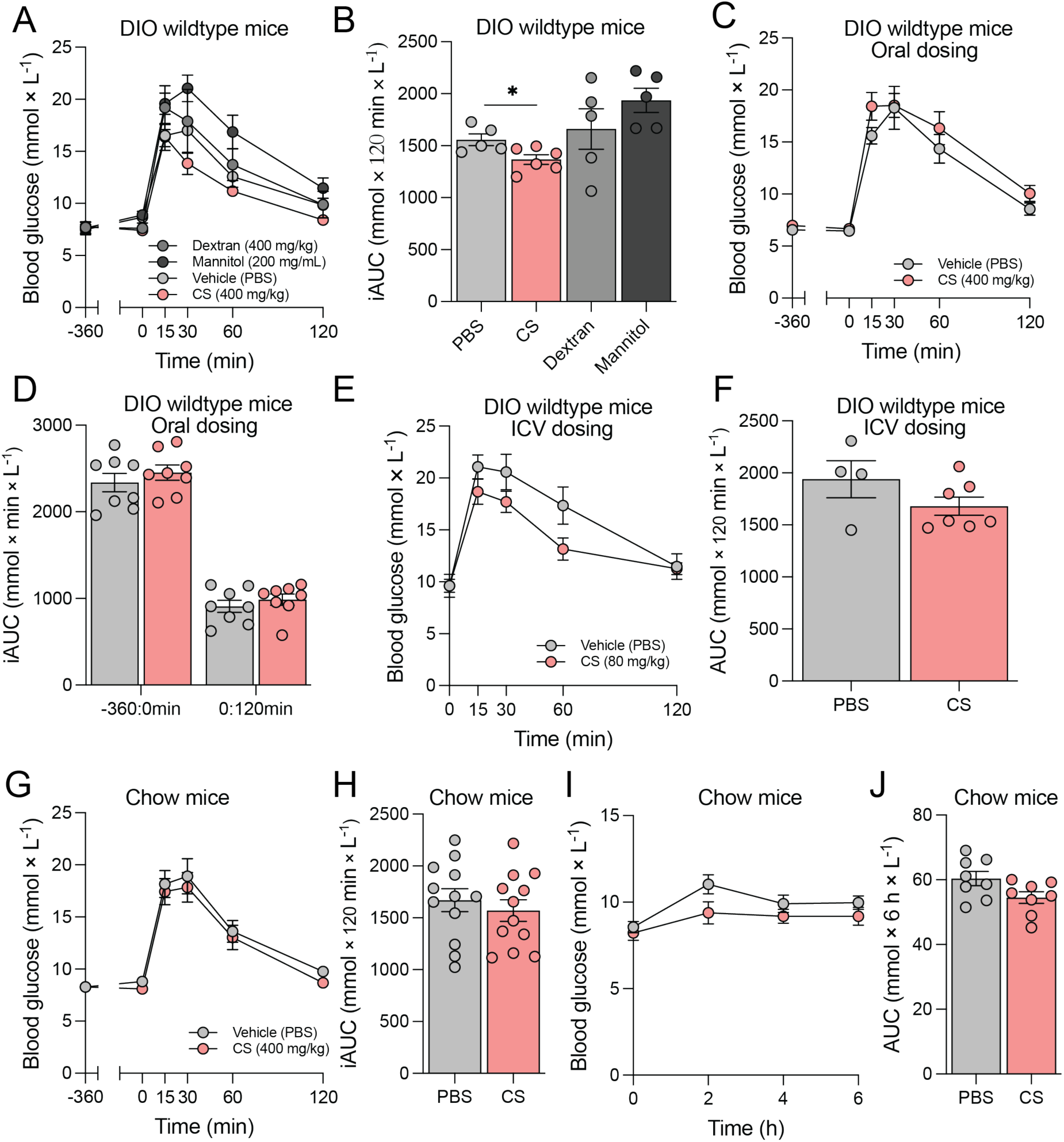
**A-B)** DIO mice were subcutaneous injected with vehicle (grey), CS (red), dextran (dark grey), or mannitol (black) 6 h before an intraperitoneal injection of glucose, and their blood glucose levels were measured (panel A) and the iAUC for 0:120 min calculated (panel B) (*n* = 5-6 per group) (mouse study 2 in Materials and Methods). The data from CS and vehicle-dosed mice are from the study presented in main figure 2E-F. **C-D)** DIO mice were administered vehicle (grey) or CS (red) by oral gavage 6 h before an intraperitoneal injection of glucose, and their blood glucose levels were measured (panel C) and their iAUCs calculated (panel D) (*n* = 8 per group) (mouse study 17 in Materials and Methods). **E-F)** DIO mice were administered vehicle (grey) or CS (red) intracerebroventricularly 4 h before an intraperitoneal injection of glucose, and their blood glucose levels were measured (panel E) and their iAUCs calculated (panel F) (*n* = 4-7 per group) (mouse study 18 in Materials and Methods). **G-H)** Chow mice were subcutaneous injected with vehicle (grey) or CS (red) 6 h before an intraperitoneal injection of glucose, and their blood glucose levels were measured (panel G) and the iAUC for 0:120 min calculated (panel H) (*n* = 5-6 per group) (mouse study 19 in Materials and Methods). **I-J)** Acute compound tolerance test of CS vs. vehicle on blood glucose levels in chow mice (panel I) and the measured AUCs (panel J) (*n* = 8 per group) (mouse study 20 in Materials and Methods). Statistics in B by one-way ANOVA. *p* value < 0.05; *. Related to main figure 2.

**Figure S4.**
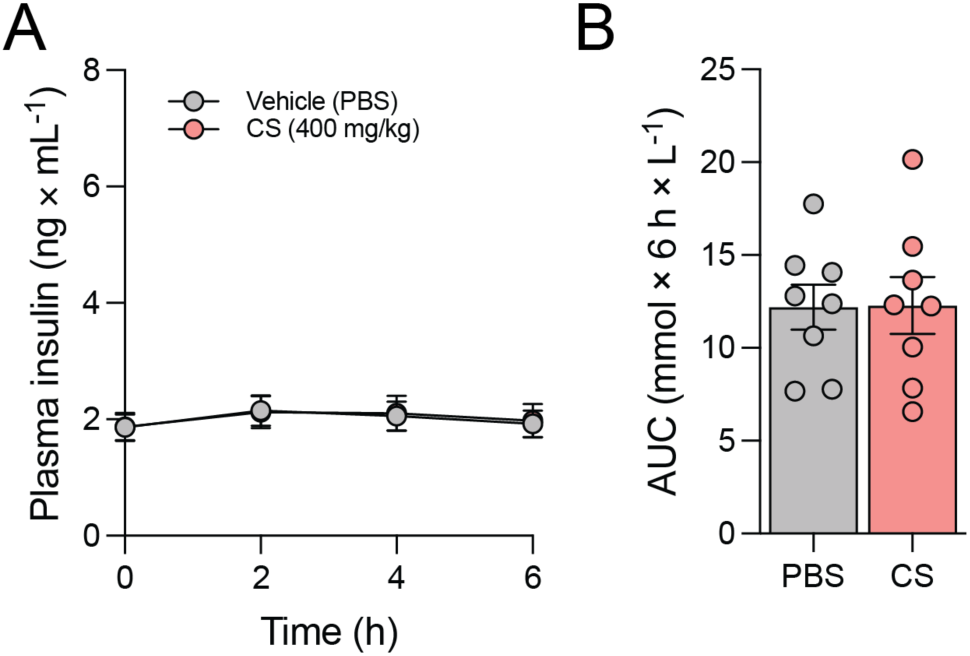
**A-B)** Acute compound tolerance test of CS vs. vehicle on plasma insulin levels in chow mice (panel J) and the calculated AUCs (panel B) (*n* = 8 per group) (mouse study 20 in Materials and Methods). Related to main figure 3.

**Figure S5.**
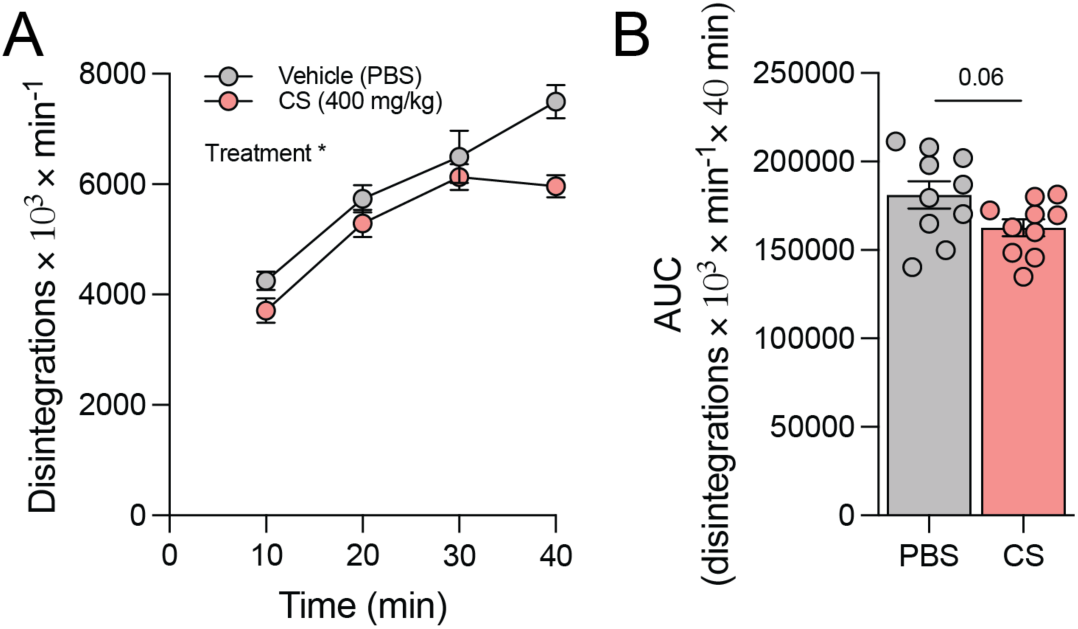
**A-B)** Glucose-stimulated glucose clearance study dependent on vehicle (grey) or CS (red) treatment (*n* = 10 per group) 6 h before an intraperitoneal injection of glucose with trace amounts of ^3^H-2-deoxy glucose (^3^H-2-DG) (mouse study 15 in Materials and Methods). Disintegrations per minute from the tracer measured from the blood (panel A) and its AUCs (panel B). Related to main figure 4.

**Figure S6.**
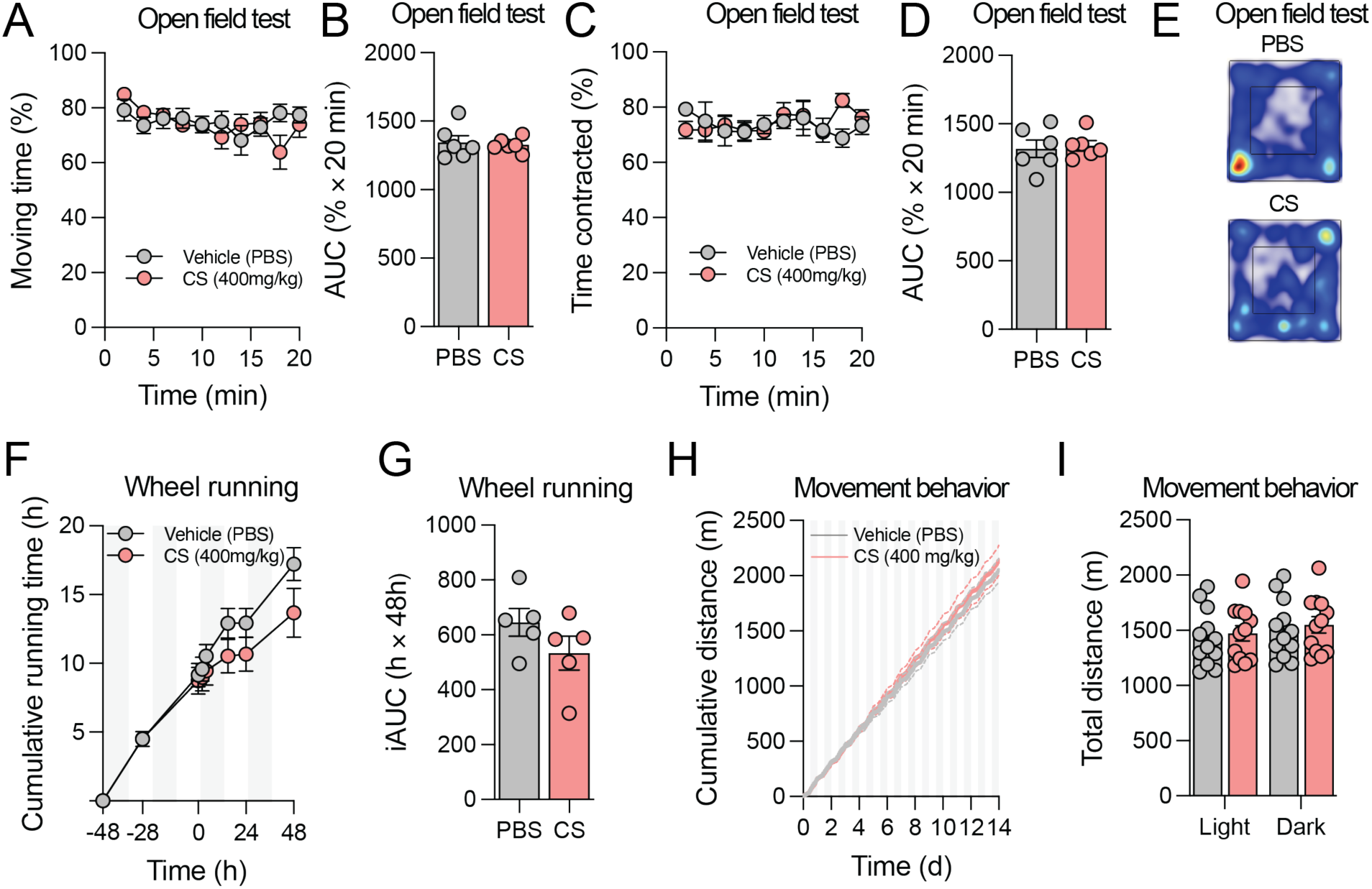
**A-E)** Open field test. Chow mice were subcutaneous injected with either CS (red) or vehicle (grey) (PBS) (*n* = 6 per group) and their behavior was followed acutely in an open field for 20 min. Moving time in percentage (panel A) and its AUC (panel B), time contracted in percentage (panel C) and its AUC (panel D) were recorded or measured in intervals as indicated. Representative heatmap traces from the open field arena provided in panel E (mouse study 7 in Materials and Methods). **F-G)** Voluntary wheel running test. Chow mice were subcutaneous injected with either CS (red) or vehicle (grey) (*n* = 5 per group) and their cumulative running time (panel F) and its iAUC for the period after injection (panel G) were measured (mouse study 8 in Materials and Methods). **H-I)** Movement behavior. DIO mice were subcutaneous injected daily with either CS or vehicle (*n* = 12 per group) in metabolic chambers and their cumulative distance recorded (panel H) and total distance calculated based on light or dark phases, as indicated (panel I) (mouse study 9 in Materials and Methods). Related to main figure 5.

**Figure S7.**
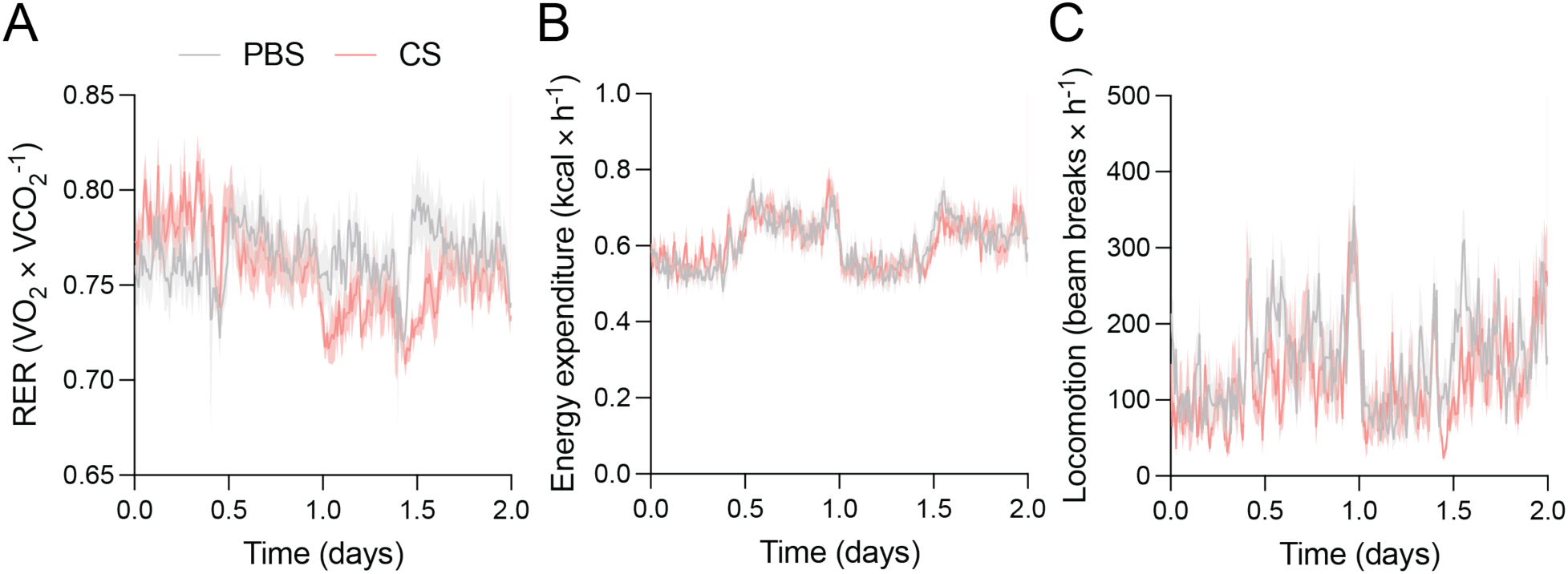
**A-C)** DIO mice (*n* = 8 per group) were subcutaneous injected with vehicle (grey) (PBS) or CS (red) daily for 2 days in metabolic chambers (mouse study 21 in Materials and Methods). Changes in respiratory exchange ratios (panel A), energy expenditures (panel B), and locomotion (panel C). Data presented as means +/-SEM and indi-vidual data points represent data from individual animals. Related to main figure 5.

**Figure S8.**
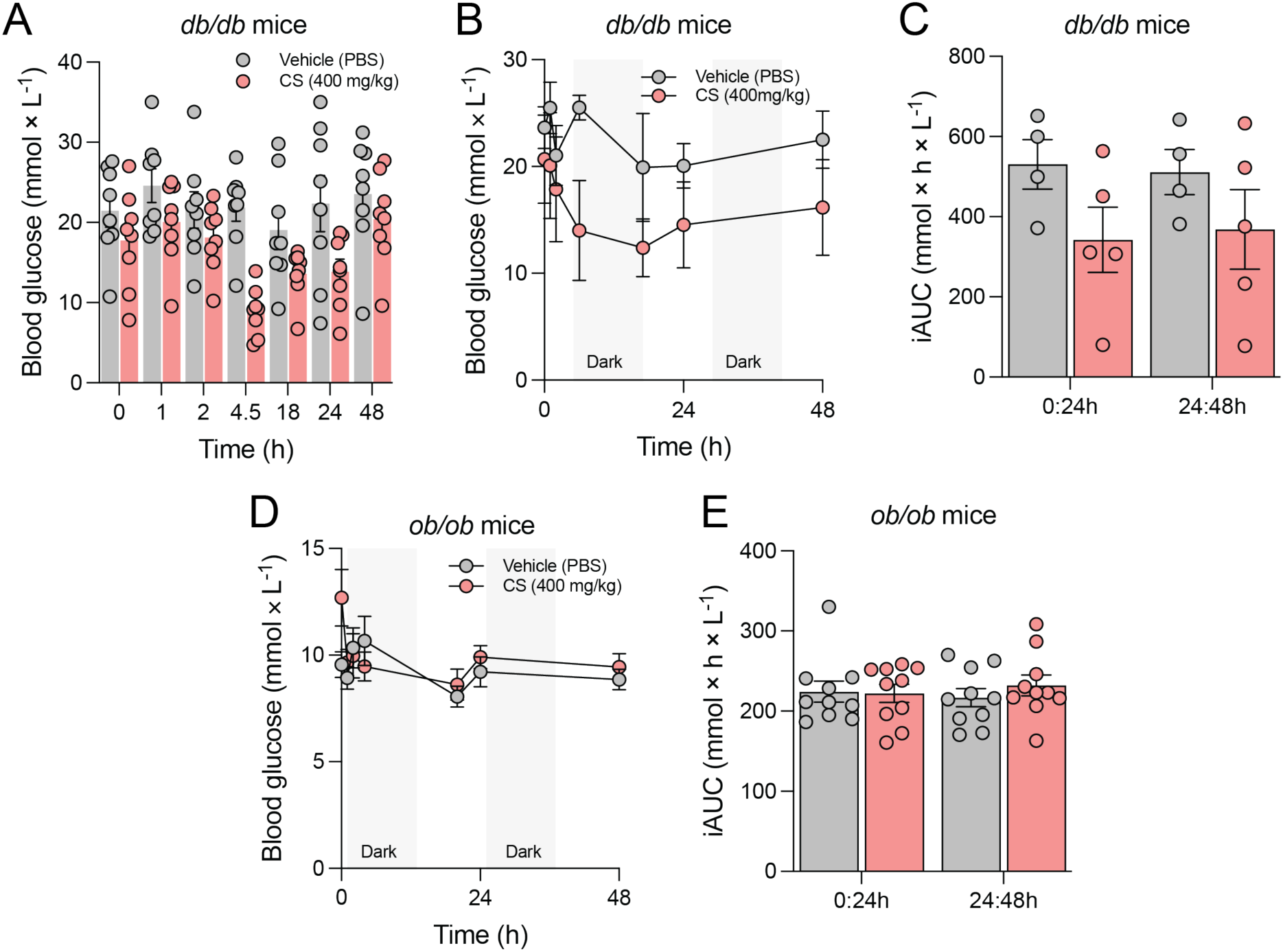
**A)** *db^-^/db^-^* mice (*n* = 8 per group) were subcutaneous injected with vehicle (grey) (PBS) or CS (red) and their blood glucose levels were measured throughout 48 h and their individual values are shown (mouse study 12 in Materials and Methods). **B-C)** Pilot study with *db^-^/db^-^* mice (*n* = 4-5 per group) where animals were subcutaneous injected with vehicle (grey) (PBS) or CS (red) and their blood glucose levels were measured throughout 48 h (panel B) and their iAUCs measured (panel C) (mouse study 13 in Materials and Methods). **D-E)** *ob^-^/ob^-^* mice (*n* = 10 per group) were subcutaneous injected with vehicle (grey) (PBS) or CS (red) and their blood glucose levels were measured throughout 48 h (panel D) and their iAUCs measured (panel E) (mouse study 14 in Materials and Methods). Related to main figure 7.

**Supplementary Table 1.**

| **Gene** | **Gene name** | **Function** | **Pathway** |
| --- | --- | --- | --- |
| XYLT1 | Xylosyltransferase 1 | Initiation:Proteoglycan | Biosynthesis |
| XYLT2 | Xylosyltransferase 2 | Initiation:Proteoglycan | Biosynthesis |
| B4GALT7 | Beta-1,4-Galactosyltransferase 7 | Core extension:Proteoglycan | Biosynthesis |
| B3GALT6 | Beta-1,3-Galactosyltransferase 6 | Core extension:Proteoglycan | Biosynthesis |
| B3GAT3 | Beta-1,3-Glucuronyltransferase 3 | Core extension:Proteoglycan | Biosynthesis |
| CSGALNACT1 | Chondroitin Sulfate N-Acetylgalactosaminyltransferase 1 | Core extension:Proteoglycan (CS) | Biosynthesis |
| CSGALNACT2 | Chondroitin Sulfate N-Acetylgalactosaminyltransferase 2 | Core extension:Proteoglycan (CS) | Biosynthesis |
| CHPF | Chondroitin Polymerizing Factor | Core extension repeat:Proteoglycan (CS) | Biosynthesis |
| CHPF2 | Chondroitin Polymerizing Factor 2 | Core extension repeat:Proteoglycan (CS) | Biosynthesis |
| CHSY1 | Chondroitin Sulfate Synthase 1 | Core extension repeat:Proteoglycan (CS) | Biosynthesis |
| CHSY3 | Chondroitin Sulfate Synthase 3 | Core extension repeat:Proteoglycan (CS) | Biosynthesis |
| DSE | Dermatan Sulfate Epimerase | Capping:Proteoglycan (CS) | Biosynthesis |
| DSEL | Dermatan Sulfate Epimerase Like | Capping:Proteoglycan (CS) | Biosynthesis |
| CHST10 | Carbohydrate Sulfotransferase 10 | Capping Sulfo:Proteoglycan | Sulfation |
| CHST3 | Carbohydrate Sulfotransferase 3 | Capping Sulfo:Wide specificity | Sulfation |
| CHST7 | Carbohydrate Sulfotransferase 7 | Capping Sulfo:Wide specificity | Sulfation |
| CHST11 | Carbohydrate Sulfotransferase 11 | Capping Sulfo:Proteoglycan (CS) | Sulfation |
| CHST12 | Carbohydrate Sulfotransferase 12 | Capping Sulfo:Proteoglycan (CS) | Sulfation |
| CHST13 | Carbohydrate Sulfotransferase 13 | Capping Sulfo:Proteoglycan (CS) | Sulfation |
| CHST14 | Carbohydrate Sulfotransferase 14 | Capping Sulfo:Proteoglycan (CS) | Sulfation |
| CHST15 | Carbohydrate Sulfotransferase 15 | Capping Sulfo:Proteoglycan (CS) | Sulfation |
| UST | Uronyl 2-Sulfotransferase | Capping Sulfo:Proteoglycan (CS) | Sulfation |
| HYAL1 | Hyaluronidase 1 | CS hydrolysis | Degradation |
| HYAL4 | Hyaluronidase 4 | CS hydrolysis | Degradation |
| SPAM1 | Sperm Adhesion Molecule 1 | CS hydrolysis | Degradation |
| GALNS | Galactosamine (N-Acetyl)-6-Sulfatase | CS hydrolysis | Degradation |
| ARSB | Arylsulfatase B | CS hydrolysis | Degradation |
| GUSB | Glucuronidase Beta | CS hydrolysis | Degradation |

